# Genetic barcoding of individual cells links cancer evolutionary trajectories and prognostic outcomes

**DOI:** 10.64898/2025.12.01.691488

**Authors:** Candice Merle, Ivan Di Terlizzi, Mathilde Huyghe, Danae Welboren, Wenjie Sun, Cécile Conrad, Leïla Perié, Steffen Rulands, Silvia Fre

## Abstract

Intratumoural heterogeneity (ITH) reflects cancer progression, by integrating clonal dynamics, cell plasticity, and metastatic potential of malignant cells. Defining the origins and evolutionary trajectories of tumour cells is central to preventing tumour growth and dissemination, yet significant gaps in knowledge persist. Here, we used an original single cell resolution lineage tracing system to investigate how ITH underlies changes in tumour cell states, such as the acquisition of an epithelial-to-mesenchymal transition (EMT) phenotype and invasive capacity. By combining *in vivo* genetic barcoding with single-cell transcriptomics, we uncover two parallel evolutionary trajectories toward EMT: one associated with alveolar differentiation, and another characterized by gene regulatory programs activated during stress and tissue regeneration. Of interest, only the gene modules defining the regenerative trajectory were found to correlate with poor survival in breast cancer patients. Our results also provide a quantitative framework to measure cellular plasticity that enables the identification of highly plastic cells and their gene signatures, correlating with rapid changes in transcriptional state, EMT features and aggressive cancer. These findings deliver a comprehensive view of *in vivo* clonal evolution in breast cancer, uncovering the interplay between lineage plasticity and tumour progression with quantitative and robust statistical approaches.

## Introduction

Cancer develops through an evolutionary process driven by the accumulation of genetic and epigenetic alterations, resulting in a mosaic of distinct, although lineage-related cells, that are defined as clones [1]. The clonal diversity of tumours, contributing to intratumoral heterogeneity (ITH), and the extensive plasticity of tumour cells synergize to promote tumour progression, but also therapy resistance and cancer recurrence. However, it remains ambiguous whether tumour growth, metastasis, and chemoresistance arise from overlapping or distinct cell subpopulations. ITH manifests at multiple levels, in the form of epigenetic modifications, genetic mutations, transcriptional changes and different cellular states, but the origins and extent of this diversity remain poorly defined [2]. As tumours progress, cells diverge from their original lineage, gaining unique traits and phenotypes that contribute to the pervasive ITH of cancers [3]. The dynamics of clonal evolution are further shaped by selective pressure from the tumour microenvironment, which influences the tumour cellular composition and fosters the adaptation and plasticity necessary for clone survival and expansion [4]. Deciphering how clones evolve at the transcriptomic level, and how cells within the same clone diverge functionally, is essential to pinpoint the cell subpopulations responsible for metastasis.

Investigating tumour evolution in human cancers presents several challenges, as clonal phylogeny is often inferred indirectly through mutations or copy number variations (CNVs) [5, 6, 7]. While this method reveals hierarchical relationships among clones, it provides limited insight into the distinct molecular features of each clone. Classical lineage tracing studies provide clonal relationships but lack molecular resolution. Due to these technical roadblocks, the connection between cell lineage and transcriptional state within native tumours remains largely unexplored. Consequently, we still do not know which molecular mechanisms are associated with clonal selection during tumour progression, and why only some tumour cells acquire EMT traits and contribute to metastases.

To address these outstanding questions, genetic barcoding has emerged as a powerful technique for tracing clonal lineage dynamics at the single cell scale and with high temporal precision [8]. When combined with mouse models for lineage tracing, barcoding enables the study of *in vivo* clonal evolution within the native tumour micro-environment. This makes it possible to determine the influence of non-heritable tumour heterogeneity, i.e. from micro-environmental cues, on clonal competition or cooperation. In this study, we employed a recently developed *in vivo* barcoding strategy that simultaneously captures clonal identity and transcriptional signature of each barcoded cell, thereby overcoming a major limitation of previous approaches. This approach enabled us to model clonal evolution in breast cancer and to reveal transcriptional signatures associated with high cellular plasticity and metastatic potential.

## Results

### Single cell RNAseq analysis of primary tumours reveals transcriptional heterogeneity

To study clonal evolution and competition *in vivo* and *in situ*, we employed the MMTV-PyMT mouse model, spontaneously developing mammary tumours that closely reproduce the features and behaviour of human luminal B breast cancers [9]. This breast cancer mouse model offers the practical advantage of rapidly developing in situ hyperplastic lesions that progress to adenomas and ductal carcinomas *in situ* (DCIS) (**Figure S1A**), which later become invasive carcinomas and eventually display lung metastases. This sequence of events occurs within a defined and highly reproducible time window of about 2 months. The hyperplasia is defined by a tissue state resembling mid pregnancy, where hyperplastic acini are filled with epithelial cells. These lesions swiftly form adenomas *in situ*, reproducing human ductal hyperplasia, characterized by extensive proliferation of epithelial cells, which are still confined by the basement membrane, and by a continuous monolayer of basal mammary cells expressing cytokeratin K14 (**Figure S1B**) [10]. At later stages, 3-month-old female mice develop carcinomas exclusively composed of luminal cells, defined by the expression of cytokeratin K8 and the loss of basal cells marked by K14. At this time, we found mixed tumours, composed of malignant nodules (referred to as “advanced” in **Figure S1A-B**) co-existing with more benign regions containing adenomas (annotated as “periphery”), highlighting the high degree of heterogeneity of these tumours [10]. To genetically label tumour cells, we chose two approaches: a first targeted technique consisted in crossing the DRAG mice to the N1Cre^*ERT2*^ mouse line, that specifically targets mammary luminal HR^*neg*^ (hormone receptor) cells (ERα-negative and PR-negative cells) [11]. A second more neutral approach used the ROSACre^*ERT2*^ mouse reporter, expressing Cre recombinase in a ubiquitous and unbiased way, that displayed GFP^*pos*^ barcoded cells in both luminal and basal mammary epithelial compartments.

When we performed single cell RNA sequencing of 6 different late-stage carcinomas derived from distinct mice, we identified three major cell types: epithelial cells, endothelial cells, and cancer-associated fibroblasts (these latter ones were categorized as “Steady-state like” SSL and vascular vCAFs, based on previous literature) [12] (**Figure S1C-E**). Uniform manifold approximation and projection (UMAP) dimensionality reduction analysis partitioned the sequenced cells into 11 clusters (**Figure 1A**). Each tumour, whether derived from the Notch1Cre^*ERT2*^ (NOTCH1-T) or RosaCre^*ERT2*^ (ROSA-T) mouse model, contributed to every cluster, thereby demonstrating the reproducibility in tumour progression in this mouse model (**Figure S1F**). Tumour epithelial cells could be further subdivided into 8 tumour epithelial cell clusters and presented a luminal signature, featuring high levels of *Epcam* and *Krt8* [13] (**Figure S1G, Figure S1G**). We scored the differentially expressed genes (DEGs) to manually annotate each UMAP cluster (**Figure 1B**). Cells in cluster 0 expressed luminal markers such as *Krt8* and *Krt18*, alongside tumour-associated genes such as *Klf5*, previously reported as an oncogene in the MMTV-PyMT model and breast cancer [14, 15]. Based on their predominant luminal signature, we termed this cluster “luminal-like cells”. Cluster 1, which we named pro-inflammatory, comprised cells with high expression of genes correlated with immune responses and inflammation such as *Csf3*, coding for the glycoprotein granulocyte colony-stimulating factor 3 and *Cxcl1*, coding for a chemokine. In contrast, cluster 2 featured strong expression of AP-1 components (*Fos, Jun*) and associated immediate early genes (*Egr1*), prompting the designation “AP1 activated”. Cluster 3 matched a luminal differentiation program toward the alveolar subtype (*Lalba, Wap* and *Scd1*), usually associated with pregnancy and lactation in the normal mammary epithelium [16]. Finally, both clusters 5 and 10 exhibited high EMT scores (**Figure S1H**).

**Figure 1:**
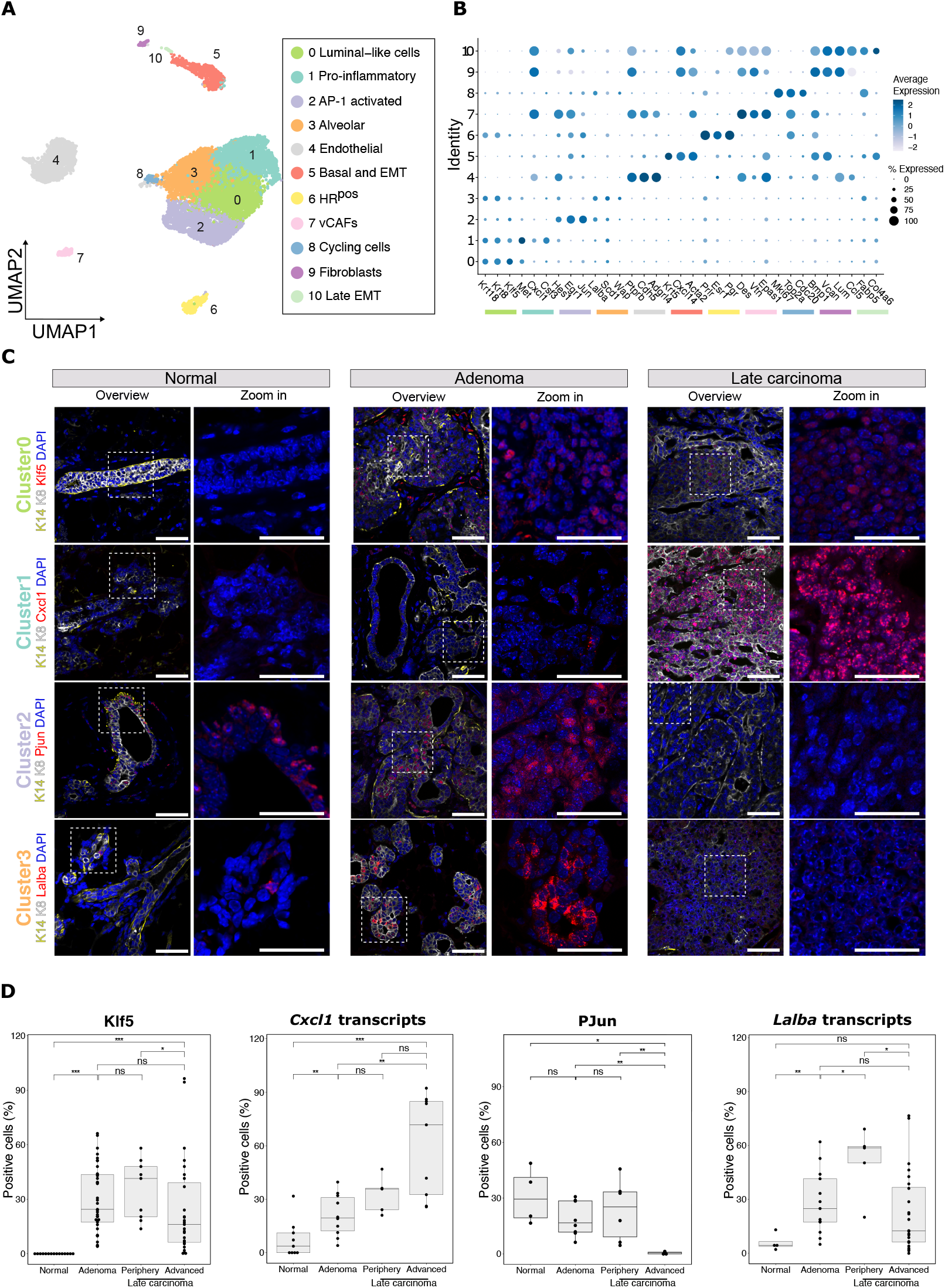
Characterization of MMTV-PyMT tumours reveals intratumoral heterogeneity and transcriptional changes during tumour progression. **A**. UMAP representation of scRNAseq analysis showing the different cell clusters identified after integration of 6 individually sequenced MMTV-PyMT late carcinomas. **B**. Dot plot representing the expression of 3 genes specific to each cluster. Dot size indicates the percentage of cells expressing each selected gene, and the colour key reflects the average expression level. **C**. Immunostainings for the main markers of tumour cell clusters in the normal mammary gland, adenomas, and late carcinomas. Scale bars represent 50 µm in overviews (left) and 25 µm in insets (right). **D**. Quantification by smRNA FISH and immunofluorescence of the indicated markers at different stages of tumour development. Each data point represents an individual region of interest (ROI) from at least n=3 tumours.

To gain further insights into the spatial distribution of these different cell clusters during tumour development and progression, we performed immunofluorescence on sections of tumours at different stages, using the markers we identified in the scRNAseq dataset. We examined marker expression across normal mammary glands, adenomas, and late-stage carcinomas. *Klf5* was expressed at early stages and maintained throughout progression to late-stage carcinoma (**Figure 1C-D**). Interestingly, we also identified markers with expression patterns characteristic of distinct stages of tumour progression. For instance, P-Jun, enriched in cluster 2 (AP-1 activated cluster), was strongly expressed in hyperplasia, but its expression dramatically decreased as tumours advanced in grade. In late carcinomas, P-Jun was restricted to residual hyperplastic regions and absent from more advanced tumour areas (**Figure 1C-D, Figure S2A**). This expression pattern was similar to *Hes1*, another signature gene of cluster 2, which was broadly expressed in adenomas but became restricted to the most differentiated regions (defined as periphery) in late-stage carcinomas (**Figure S2B-C**). Likewise, *Lalba*, enriched in cluster 3, showed dramatic enrichment in adenomas compared to the normal gland, and its expression remained stable in late carcinomas (**Figure 1C-D**). However, *Lalba* expression was heterogeneous across tumours, with some regions showing strong expression and others devoid of it (**Figure S2D**). Finally, *Cxcl1*, a specific marker of the “pro-inflammatory” cluster 1, exhibited a progressive increase in expression with tumour stage, likely reflecting its association with the most aggressive tumour cells (**Figure 1C-D**).

To further investigate early tumour development, we also performed scRNA-seq on early hyperplastic lesions (**Figure S2E-F**). This analysis revealed cell clusters corresponding to alveolar state (*Lalba* expression in clusters 2 and 3), cycling cells (*Mki67* expression in cluster 8), AP-1 pathway activation (*Jun* expression in cluster 4), luminal differentiation (clusters 0 and 1, enriched in *Krt8*), and basal/myoepithelial cells (cluster 5). Consistent with our previous observations, *Cxcl1* expression was limited to a small subset of luminal cells in hyperplasia (**Figure S2F**). This analysis confirmed that early stages of MMTV-PyMT tumour development involve both activation of the AP-1 pathway and acquisition of an alveolar-like state. Notably, the presence of a defined pro-inflammatory cluster appeared to be specific to late-stage carcinomas.

### Clonal dynamics and competition during tumour progression

Having established the transcriptional signature of each tumour cell and defined the dynamics of expression patterns during tumour progression, we sought to investigate how these heterogeneous cellular states emerge and evolve at the clonal level. First, to spatially evaluate clonal evolution in these tumours, we employed Confetti-based lineage tracing, which enables visual tracking of clonal dynamics over time in vivo. This approach allowed us to assess how the distribution of different cell clones evolved during tumour progression and to determine whether specific clones expanded or became restricted as tumours advanced to more aggressive stages. Longitudinal imaging of the ROSACre^*ERT2*^/MMTV-PyMT/Confetti mouse model revealed the coexistence of multiple distinct clones within early-stage adenomas (**Figure 2A**). As the tumour progressed, we observed a marked reduction in clonal diversity, accompanied by the increase in size of the surviving clones. This dynamic behaviour illustrates the process of clonal selection that occurs during tumour progression, whereby a few clones expand at the expense of less fit ones which are lost [17].

**Figure 2:**
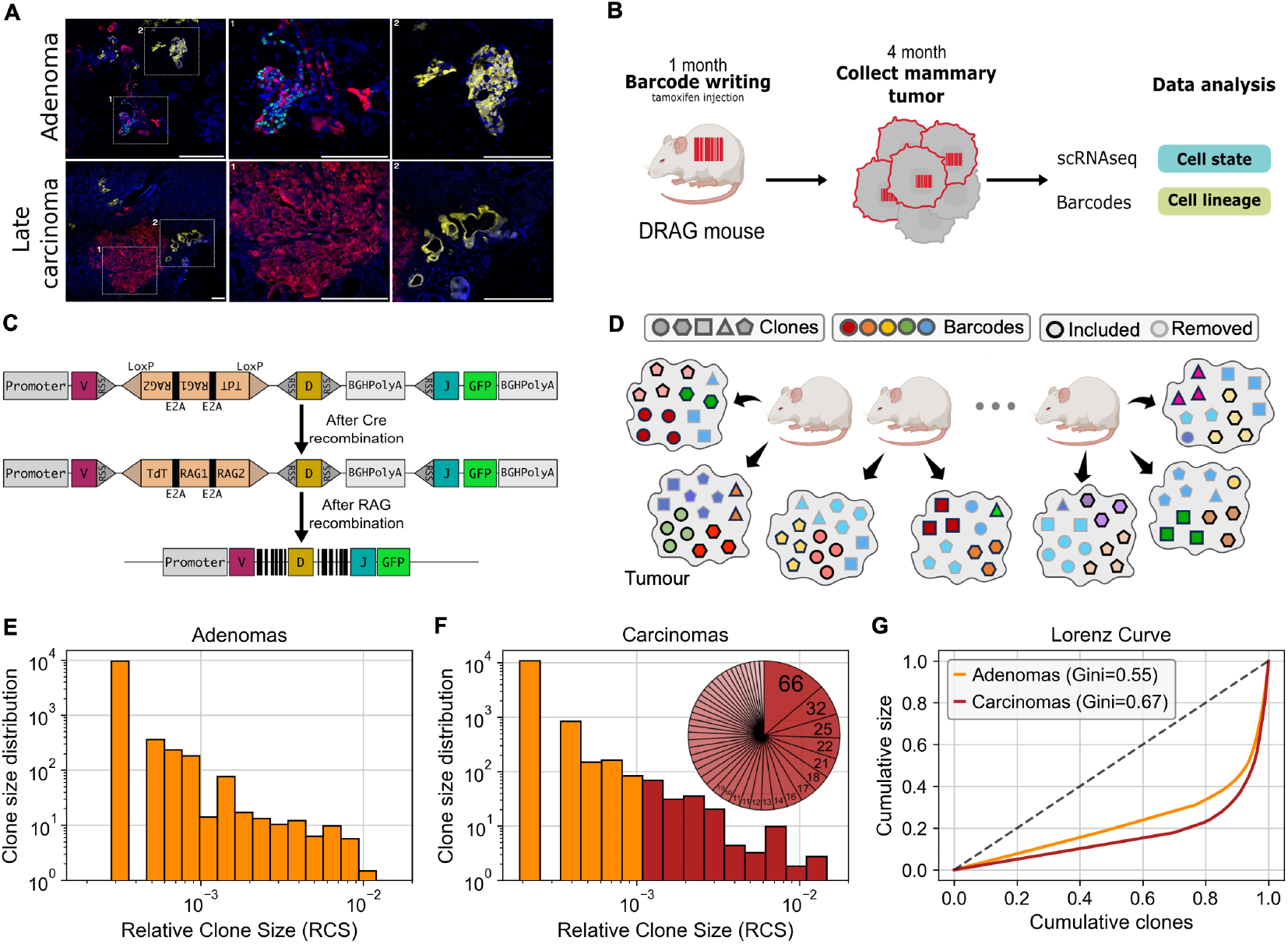
DRAG barcoding reveals clonal expansion dynamics during tumour progression. **A**. Representative images of Confetti-traced tumour cell clones in adenomas and late carcinomas. Here, clones appear as groups of cells sharing the same fluorescent colour. While both stages exhibit multicellular clones, late carcinomas display larger clones and fewer colours. **B**. Experimental workflow for barcode labelling in the MMTV-PyMT mouse model. **C**. Schematic of the original DRAG allele before and after Cre-mediated and DRAG recombination. **D**. Schematic of the barcode filtering strategy. Cells belonging to the same clone are depicted with the same shape while barcodes are colour-coded. They are subsequently evaluated and filtered based on their recurrence across tumours. In particular, barcodes found in multiple samples, are flagged as potentially non-monoclonal and removed (grey edges), while less frequent barcodes are used for downstream analysis (black edges). **E**,**F**. Clone size distributions for adenomas (C) and late carcinomas (D) after barcode filtering. A Mann–Whitney test yields a statistic of 7 · 10^5^ with *p* ≈ 5 · 10^−27^. In carcinomas, bars are colour-coded by clone size (dark orange: <5 cells; red: ≥5 cells). The inset pie chart illustrates the relative contribution of large clones (≥5 cells). **G**. Lorenz curves comparing clonal size inequality in adenomas and late carcinomas. The curves show the cumulative fraction of sample size as a function of the cumulative fraction of clones (sorted by size). Carcinomas display a stronger deviation from the diagonal compared to adenomas, reflected by higher Gini coefficients, indicating greater inequality and stronger dominance by a few large clones.

While the Confetti lineage tracing system enables visual tracking of clonal heterogeneity, it is inherently limited by the small number of fluorescent labels it can generate. To overcome this limitation and achieve high-resolution tracking of tumour clonal evolution, we employed *in vivo* genetic barcoding using the DRAG (Diversity through RAG) mouse model, generating a large repertoire of unique barcode sequences [18]. This model permits to define the identity of each barcoded cell (cell state), based on its transcriptional signature, alongside its lineage, derived from barcode based lineage tracing (cell lineage) (**Figure 2B**). This approach allowed us to trace lineages derived from barcoded cells toward the formation of distinct clones during tumour growth, while retrieving the transcriptional profile of individual clones, revealing the transcriptomic programs that drive clonal expansion and cell plasticity. In our system, barcode diversity arises through stochastic V(D)J recombination, a process well-characterized in the adaptive immune system for generating T-cell receptor gene rearrangements (**Figure 2C**) [19]. Barcode writing and expression is coupled with GFP expression, allowing FACS sorting of barcoded cells and their in vivo lineage tracing (**Figure S2G-H**). The DRAG mouse model uses this inherent variability to create a potentially unlimited number of genetic barcodes, ensuring an ample supply of unique identifiers to monitoring clonal dynamics across several tumour samples [18]. Although barcodes are generated in large numbers, their rearrangements occur at highly unequal frequencies, and some barcodes arise multiple times within the same sample. These non-unique identifiers, which we term “popular barcodes”, can be independently generated in distinct cells of origin that are not part of the same clone (e.g., barcodes shown in different shades of blue in **Figure 2D**). To mitigate this drawback complicating the assignment of the clonal origin of each barcoded cell, we developed a statistical method to estimate the probability that a barcode is truly monoclonal, i.e. cells labelled with a given barcode belong to one single clone within a tumour. For each barcode, we calculated this probability based on its frequency across samples, thereby estimating the likelihood that it originated from a single unique clone. More specifically, our method calculates the posterior probability, *p*_MC_, that a barcode, observed a given N number of times across all samples, is truly monoclonal (see the Methods Section for mathematical details). This probability of monoclonality is inversely proportional to the frequency of detection of the same barcode, as a high occurrence suggests the barcode may span multiple clones. By setting a threshold (*p*_MC_ ≥ 0.9 in our case), we removed barcodes with a low probability of being monoclonal and retained only those featuring a high statistical confidence of representing individual clones.

Following this filtering step, we examined the distribution of clonal sizes at two stages of tumour development: adenoma and late carcinoma. To account for differences in sample size, we represented clonal sizes with the relative clone size (RCS), defined as the proportion of tumour cells belonging to a given clone (**Figure 2E-F**). Both tumour stages exhibited highly skewed distributions, where most clones contributed minimally, while a few clones accounted for a substantial fraction of the tumour mass. To test whether this imbalance was more pronounced in late carcinomas, we first compared the two distributions using a Mann–Whitney test, which yielded a test statistic of 7 · 10^5^ and a p-value of approximately 5 · 10^−27^, confirming a highly significant difference between the clonal distribution in adenomas and carcinomas. We then summarized these distributions using Lorenz curves and the corresponding Gini coefficients (**Figure 2G**). The Lorenz curve describes inequality, i.e. if only few clones contribute to the majority of the tumour mass, by plotting the cumulative fraction of tumour mass (y-axis) against the cumulative fraction of clones (x-axis), after ranking them by size. The diagonal corresponds to a perfectly equal scenario, in which all clones contribute equally to the tumour. Deviations below this diagonal reveal unequal contributions, with stronger curvature indicating that a few clones dominate. The Gini coefficient provides a single quantitative measure of this inequality, with values ranging from 0 (all clones are equipotent) to 1 (maximum inequality, where one single dominant clone contributes to the entire tumour). Adenomas had a Gini coefficient of 0.55, while carcinomas reached 0.67, indicating a marked increase in clonal dominance as tumours progress.

To summarise, this statistical framework allowed for a rigorous and statistically robust assessment of barcode reliability, enabling an unbiased analysis of the tumour clonal structure and cellular hierarchy. By combining barcode filtering with quantitative measures, we found that clonal dominance increases significantly during tumour progression. Although early adenomas already show an imbalanced clonal size distribution, this imbalance becomes more pronounced in late carcinomas, where a few expanded clones dominate the population.

### Distinct evolutionary trajectories associated with the EMT state

Building on our observations that clonal distribution follows distinct growth modes across tumour stages, we next investigated whether the progression toward EMT occurs through a common transcriptional program or via multiple, clonally distinct trajectories. To distinguish between these scenarios, we analysed how individual clones, identified by their unique barcode sequences, were distributed across transcriptional states (UMAP clusters). For each clone, we quantified the proportion of cells assigned to a given UMAP cluster, generating a vector that quantitatively captures the transcriptional heterogeneity of each clone. Then, we measured the differences in distribution among UMAP clusters between clones, using Cosine similarity (see Methods Section for details), where a value of 0 indicated identical distributions across transcriptional states of cells carrying the same barcode, while values closer to 1 indicated increasingly divergent transcriptional state occupancy, with clonal lineages spanning across many different UMAP clusters. Hierarchical clustering based on the cosine distance classified the clones into eight major groups, which we refer to as *Fate Clusters* (FC). Each Fate Cluster comprises several clones that share similar distributions across transcriptomic states (**Figure S3A**). As an internal validation, we found that Fate Cluster 1 exclusively comprises several clones composed of cells from the endothelial cluster, which are not expected to have lineage relationship with epithelial tumour cells (**Figure 3A-B, Figure S3A**).

**Figure 3:**
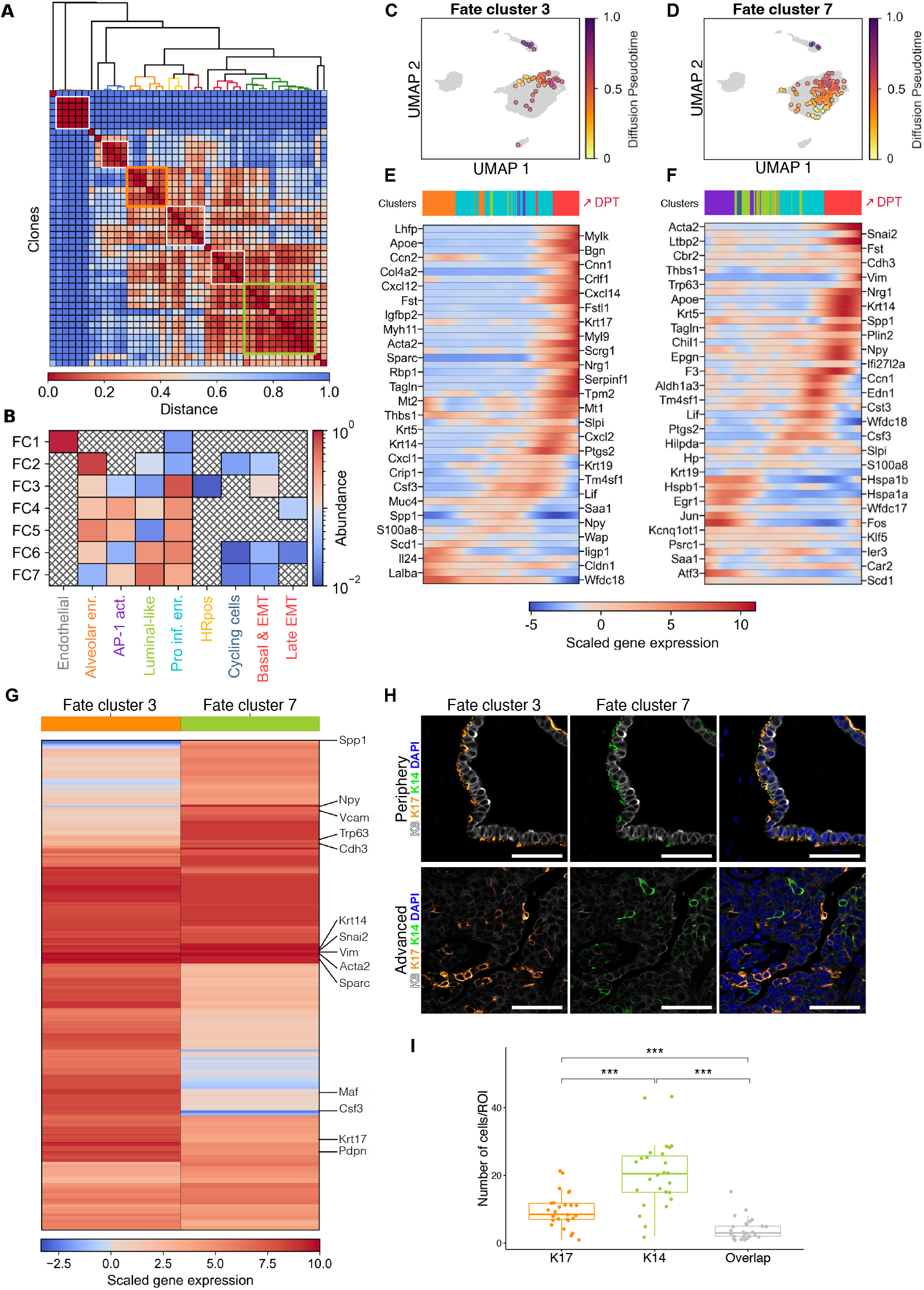
Lineage-informed transcriptomic analysis reveals distinct clonal trajectories toward EMT. **A**. Hierarchical clustering of clones using cosine similarity calculated on the distribution of each clone across transcriptional states. **B**. Heatmap showing the percentage of cells in each transcriptional state (UMAP cluster) for each Fate Cluster. Fate Cluster 1 contains exclusively endothelial cells, confirming classification specificity. **C, D**. UMAP projections of cells assigned to Fate Clusters 3 and 7, respectively. Cells are coloured by diffusion pseudotime (colour key shown). **E, F**. Gene expression dynamics along diffusion pseudotime for Fate Clusters 3 (E) and 7 (F). **G**. Heatmap of EMT-associated marker expression accross cells from Fate Clusters 3 and 7. **H**. Immunofluorescence on MMTV-PyMT late-stage carcinoma showing mutually exclusive expression of markers specific to Fate Cluster 3 (K17) and Fate Cluster 7 (K14). **I**. Quantification by immunofluorescence of the cells expressing either K14 or K17 or both in late stage carcinomas. Data points represent an individual region of interest (ROI). Scale bars represent 50µm.

To understand how clones transition toward the EMT state, we focused on Fate Clusters containing EMT-marked cells. Two dominant patterns emerged. Fate Cluster 3 included clones that spanned alveolar, pro-inflammatory and EMT clusters (**Figure 3B, Figure S3A**). Fate Cluster 7, instead, connected the AP-1 activated, luminal-like, pro-inflammatory and EMT clusters, creating two distinct and parallel transcriptomic profiles leading to EMT. In addition, Fate Cluster 6 contained mixed clones spanning luminal-like, pro-inflammatory, alveolar, and EMT states, indicating the potential existence of a more complex or plastic transition for some cells. To investigate the transcriptional dynamics underlying these molecular profiles, we applied pseudotime analysis [20] to Fate Clusters 3 and 7, to order cells along a transcriptional axis. Pseudotime provides a continuous measure of progression that arranges cells according to their transcriptional similarity, thereby reconstructing dynamic processes from static single-cell data. In this analysis, the EMT phenotype emerged as the terminal state of the trajectory (**Figure 3C-D**). While both trajectories implied passage through an intermediate inflammatory state, they originated from mutually exclusive cell states, either the alveolar or the AP-1 activated cluster, suggesting that different clonal lineages progress toward EMT via separate, although concomitant, transcriptional programs. For Fate Cluster 3, pseudotime analysis revealed a transition from alveolar cells expressing alveolar luminal differentiation markers (*Krt19, Wap, Lalba*), to pro-inflammatory states characterized by upregulation of cytokines such as *Cxcl1* or *Csf3* or interleukine *Il24*, and terminating with the EMT-associated expression of *Scrg1* and *Apoe* (**Figure 3E**). For Fate Cluster 7, the trajectory began with a high expression of stress-response genes (*Egr1, Fos* and *Jun*), which then acquired the expression of *Klf5*, and progressively started to express genes related to basal cells such as *Krt14, Acta2* and invasive markers, such as *Vim, Cdh3* and *Snai2* (**Figure 3F**). Of note, these transcriptional trajectories were not observed in hyperplastic lesions at early stages of tumorigenesis, suggesting that they arise later in tumour evolution.

To assess whether the two Fate Clusters leading to EMT via alternative routes converged to a similar endpoint state, we then analysed the gene expression profiles associated with the EMT state in both clusters. EMT states associated with Fate Cluster 3 and 7 shared the expression of common genes such as *Acta2, Apoe* and *Tagln*, all associated with an EMT phenotype. This comparison revealed also that distinct gene signatures characterize these two different EMT states (**Figure 3G**). The EMT state associated with Fate Cluster 3 featured the expression of genes associate with an immune response, such as *Cxcl1, Cav1* and *Cxcl14*, related to GO terms such as “positive regulation of defence response” (**Figure S3B**). The EMT phenotype originating from Fate Cluster 7, instead, was enriched in expression of genes associated with migration (*Vim, Snai2* and *Klf4*) and wound healing (*Mylk, Ptn* and *Serpine1*). This analysis demonstrates that the two Fate Clusters lead to distinct EMT states: one associated with an immune response while the other is characterized by an invasive and regenerative phenotype. Of interest, EMT markers from these two trajectories show non-overlapping localization in cells of late-stage carcinomas, even though they are co-expressed in basal mammary cells of normal glands (**Figure 3H-I**).

### Distinct clonal evolutionary programs during cancer progression harbour prognostic implications

Having identified the genes associated with the two clonal trajectories leading to different EMT states, we next sought to characterize the transcriptional programs underlying these clonal evolution paths. Rather than relying solely on differential gene expression analysis, which typically focuses on pairwise comparisons and can oversight broader co-regulated patterns, we used the clonal information revealed by our barcoding strategy to associate gene expression programs with specific clonal evolutionary routes (see Methods Section). Using this approach, we could identify gene modules whose expression patterns aligned with distinct clonal trajectories, offering insights into the transcriptional programs underlying divergent clonal evolution. We then applied the Hotspot method [21] to classify groups of genes (that we named Gene Modules) that co-vary across cells within each Fate Cluster and obtained 14 robust Gene Modules (**Figure 4A**).

**Figure 4:**
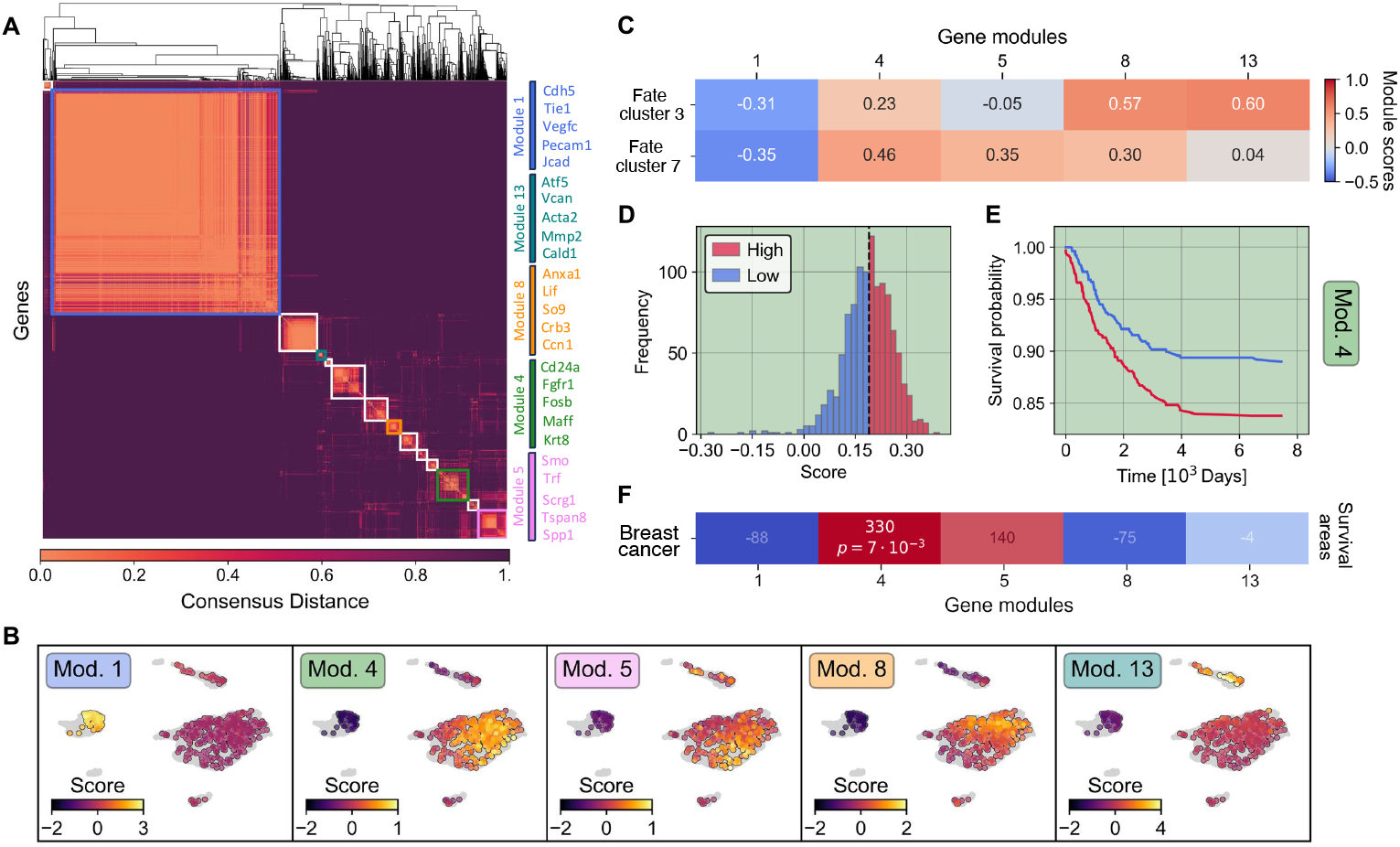
Distinct routes of clonal evolution are associated with different clinical outcomes in human tumours. **A**. Gene modules identified by Hotspot, organized by consensus clustering across Fate Clusters. **B**. UMAP representations showing expression scores for the selected gene modules. **C**. Heatmap displaying the average Gene Module Scores for the indicated modules across Fate Clusters 3 and 7. **D**. Histograms of Gene Module Scores for module 4, associated with Fate Cluster 7, in the BRCA (Breast Invasive Carcinoma) TCGA dataset. **E**. Kaplan-Meier survival curves for gene module 4 in human breast cancer (BRCA from TCGA). **F**. Heatmap showing the area between survival curves associated to selected gene modules for breast cancer patients. Survival differences not reaching statistical significance are blurred; significant values are displayed along with the corrected *p*-values.

To correlate Gene Modules and Fate Clusters, we computed a score, termed Gene Module Score, for each Gene Module in every Fate Cluster, that takes into account the average expression of the genes belonging to each module. A higher Module Score indicates that the genes in the module are expressed at higher levels in that Fate Cluster than would be expected by chance (see Methods Section for details). Therefore, Module Scores quantify the extent to which a specific gene program is upregulated across different clones and lineage trajectories. To illustrate the functional relevance of this approach, Gene Module 1 was highly associated with Fate Cluster 1, representing only endothelial clones, and included genes such as *Cdh5, Pecam1* and *Plpp3*, linked to the “endothelial cell migration” GO term (**Figure S4**).

We then focused on Gene Modules associated with the two main clonal trajectories leading to EMT. Fate Cluster 3 showed high scores for Gene Modules 8 and 13 (**Figure 4B-C**). These Gene Modules are linked to developmental processes (e.g., *Ccn1, Cebpb*) and extracellular matrix organization (e.g., *Timp2, Fermt1*)(**Figure S4**). In contrast, Fate Cluster 7 was predominantly associated with Gene Module 4, which includes genes such as *Cd24a, Fgfr1*, and *Muc1*, linked with morphogenesis and with Gene Module 5, including genes expressed in response to stress. These findings reveal that the two EMT trajectories are driven by exclusive gene programs.

Since Fate Clusters 3 and 7 led to two different EMT states and were also associated with different transcriptomic programs, we next asked if they were associated with different clinical outcomes. To answer this, we interrogated The Cancer Genome Atlas program (TCGA dataset) to assess the predictive power of patient survival for the identified Gene Modules in human breast cancer [22]. Intriguingly, high expression of the genes associated with Gene Module 4 and correlated with the Fate Cluster 7 was significantly predictive of poor prognosis in breast cancer patients, suggesting that this pathway may be linked to aggressive disease (**Figure 4D-E**). In contrast, the Gene Modules associated with Fate Cluster 3 showed no statistically significant association with patient prognosis in human breast cancer. These results suggest that the two routes of EMT are not only molecularly distinct but also associated with different clinical outcomes. In our mouse breast cancer model, two clonal transcriptomic programs led to EMT, but only the trajectory associated with a regenerative signature was linked to poor prognosis in human patients, supporting the idea that not all EMT programs have equal capacity of driving metastasis in breast cancer.

### Quantitative and lineage-based analysis of cellular plasticity

The distinct EMT programs that we identified and characterized reflect different transcriptional landscapes and highlight clonal differences that contribute to ITH. With the results obtained, we could then explore another key driver of tumour progression, cellular plasticity, at single cell resolution within each clone. Cellular plasticity, that we defined as the ability of cells to adapt to diverse environments and to transition between cell states, plays a crucial role in tumour development and cell heterogeneity. In an evolutionary context, plasticity enables single cells within the same cluster to acquire a broad range of phenotypic states, thus increasing the likelihood of survival under changing or adverse conditions. Indeed, plasticity can confer a survival advantage by facilitating transitions into cell states better suited to evade immune attacks, to resist treatment, or to colonize new niches. Within each clone, we identified cells that had reached the EMT state as well as cells at earlier points along the trajectory, demonstrating varying degrees of cellular plasticity within one same clone. To reveal genes and transcriptional programs associated with plasticity in the MMTV-PyMT mouse model, we first identified highly plastic cells within each barcoded clone. Cellular plasticity was inferred within a given clone and reflected the capacity of some cells to adopt different transcriptional states compared to other cells within the same clone. To quantify this, we calculated a plasticity score that integrates the number of transcriptional states found within a clone and their position along the pseudotime toward EMT. Starting from the cell with lowest pseudotime in the clone, the plasticity score increased whenever a cell positioned later in pseudotime was found in a new, never experienced transcriptional state (as defined by the UMAP clusters in **Figure 1A**). The final score for each cell was calculated as the product of this cumulative count of unique states and the cell pseudotime (see Methods Section for details). This design captures an essential aspect of plasticity: clones that spread across multiple states and extend further along the pseudotime receive higher scores, reflecting their broader transcriptional repertoire and temporal progression. This computational method implies that some larger clones composed of cells that explore more transcriptional states also tend to receive higher plasticity scores, an expected conclusion given that these clones are more likely to exhibit adaptability and fitness advantages. However, we also observed large clones restricted to a single transcriptional state and thus featuring a relatively low plasticity score. Using this metric, we classified high-plasticity cells those within the top 10% of the score distribution and low-plasticity cells those falling below this threshold (**Figure 5A–B**).

**Figure 5:**
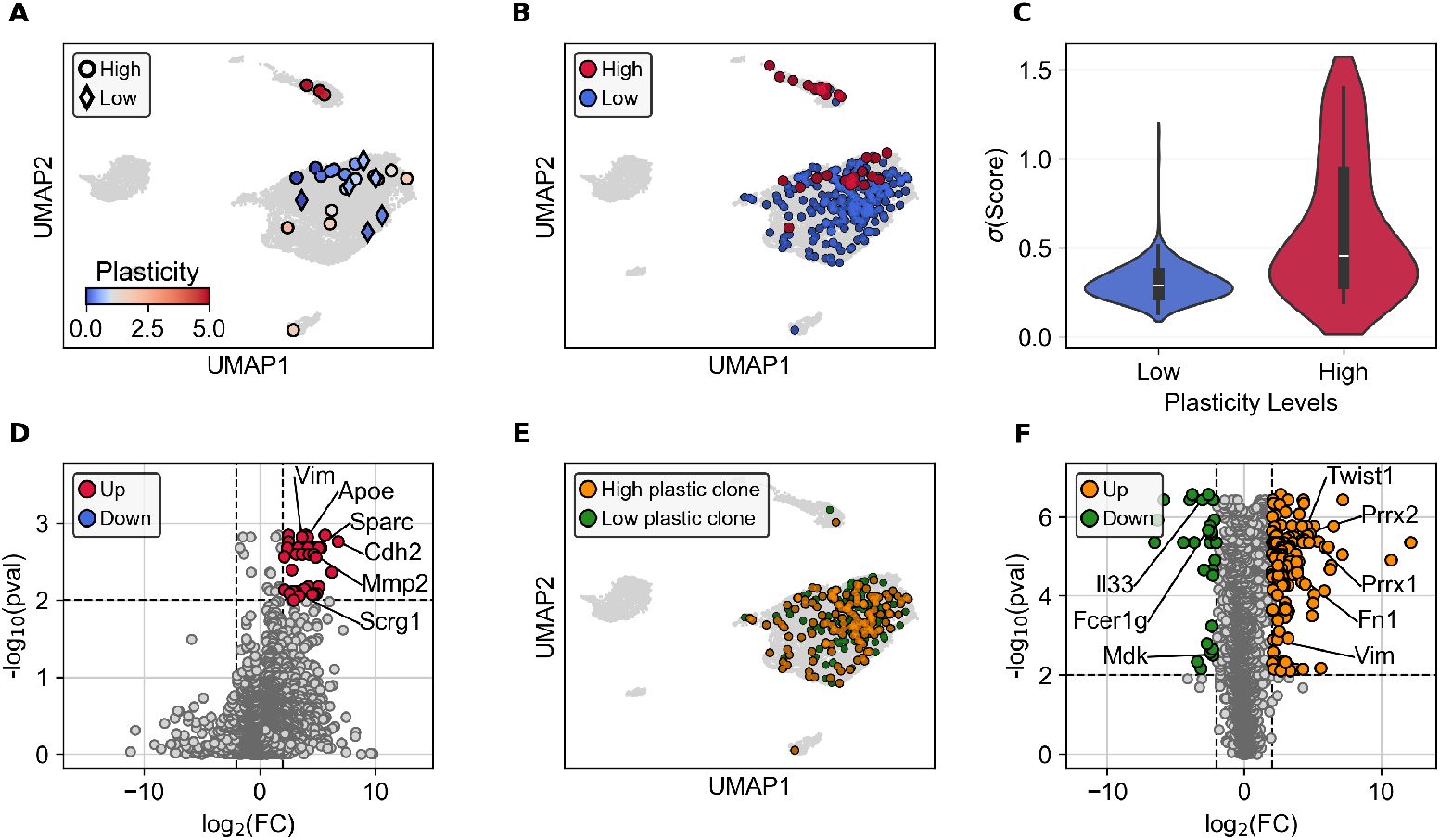
Highly plastic cells share a common transcriptional signature. **A**. UMAP showing example of 2 clones composed exclusively of low-plasticity cells (diamonds) or spanning a broad range of plasticity values (circles). Each dot represents a single cell and is coloured by its plasticity score. **B**. Distribution of high-(red) and low-plasticity (blue) cells across the transcriptomic landscape. **C**. Violin plots showing the distribution of the standard deviations of Gene Module scores for high (red) and low (blue) plasticity cells. Highly plastic cells consistently exhibit greater score variability, indicating a more heterogeneous transcriptional state rather than uniform upregulation of all programs. **D**. Volcano plot showing genes enriched in highly plastic (red) versus low-plastic (blue) cells. **E**. UMAP showing low-plasticity cells from clones that also contain highly plastic cells (orange, “high plastic clones”) or from clones composed entirely of low-plasticity cells (green, “low plastic clones”). **F**. Volcano plot showing genes differentially expressed between low-plasticity cells from high-plastic clones (orange) and those from low plastic clones (green).

We next examined how cell plasticity relates to transcriptional heterogeneity by comparing the variability of Gene Modules (from our previously described Hotspot analysis) across cells (**Figure 5C**). High plasticity cells displayed significantly higher gene expression variability, suggesting that they do not uniformly activate transcriptional programs but instead flexibly engage the expression of distinct transcriptional program simultaneously. This flexible regulation may support transitions between states and increase adaptability to different tumour niches. Similar patterns of transcriptional variability have been associated with stemness and multilineage potential in normal tissues [23, 24].

To further understand the molecular underpinnings of cell plasticity, we performed a differential analysis of gene expression between high- and low-plasticity cells, identifying genes with significant changes (**Figure 5D**). Although no genes were significantly associated with low plasticity, highly plastic cells exhibited enrichment of several genes associated with EMT, including *Vim, Apoe, Scrg1* and *Cdh2*, and annotated with the GO terms “tissue migration” and “extracellular organisation” (**Figure S5**).

Some clones were composed of cells acquiring different transcriptomics states, whereas other clones never gave rise to cells with high plasticity. To address whether plasticity emerges in specific cells or reflects a broader clonal potential, we next focused on the low-plasticity cells, and compared those originating from homogeneous clones (composed entirely of low-plasticity cells) to those derived from heterogeneous clones (containing both low- and high-plasticity cells) (**Figure 5E**). Although similar in terms of plasticity score, low-plasticity cells from heterogeneous clones displayed a distinct transcriptional signature enriched for EMT-related genes such as *Twist1, Prrx1/2, Fn1*, and *Vim* (**Figure 5F**). This indicates that even before acquiring high plasticity, these cells exhibit a primed transcriptional state predictive of future transitions, suggesting that plasticity is a clonal property encoded in the cell state long before phenotypic diversification becomes evident.

## Discussion

Understanding how clonal diversity arises and contributes to ITH remains a major challenge in cancer research. Existing approaches often capture either the lineage information or the transcriptional state of a given cell, but rarely both at once, limiting our ability to reconstruct clonal dynamics in vivo within the complex tumour ecosystem. In this study, we simultaneously captured clonal relationships and transcriptional states of single cells across different tumour stages. This approach revealed distinct evolutionary trajectories with molecular signatures associated with specific tumour clones, shaping ITH and tumour progression. Unexpectedly and of medical relevance, these clonal paths also correlated with differential prognosis in human breast cancer, demonstrating a link between clonal dynamics and tumour cell aggressivity and plasticity.

We found that transcriptional heterogeneity increased with the tumour stage, determined by changes in clonal diversity. The most aggressive clonal trajectory originates from a progenitor-like state and progresses toward a mesenchymal, regeneration-related EMT phenotype. A progenitor state associated with an injury phenotype has been described during squamous cell carcinomas and colorectal cancer progression [25, 26]. This cellular trajectory mirrors injury-repair programs observed in other cancers and reflects a high degree of cellular plasticity and adaptability, features often linked to therapy resistance and metastatic potential. Tumours coopt regenerative programs involving upregulation of stress signalling, often mediated by AP-1 activation, thereby promoting cancer cell plasticity. Dysregulation of these transcriptional programs generates highly adaptable cell clusters, associated with lineage infidelity and high plasticity, capable of tumour growth, immune evasion and metastasis.

Interestingly, we found that only the gene modules associated with the regenerative trajectory (Fate Cluster 7) were significantly correlated with poor prognosis in breast cancer, in agreement with the literature [27]. Conversely, the EMT program linked to immune activation did not associate with poor outcomes, suggesting functional diversity among EMT states. These two clonal trajectories are also associated with distinct EMT end points [27, 28, 29]. This diversity is likely influenced by microenvironmental factors, such as proximity to blood vessels or immune interactions, or by different cells of origin [29]. However, a recent study also identified two trajectories in cancer and reported that perturbing markers of one EMT trajectory can trigger the expression of genes associated with the other trajectory, highlighting the elevated intrinsic plasticity of cancer cells, independent of the acquisition of novel mutations or environmental cues [27]. This highlights the importance of investigating not only the different transcriptomic states or EMT programs, but also the plasticity of cells that drives changes in transcriptional states during tumour progression, to have a comprehensive understanding of ITH.

It is also interesting to note that, beyond interclonal heterogeneity, we also observed intraclonal differences: within a single clone, sharing the same barcode, some cells reached an EMT state while others did not and trailed behind in pseudotime. Our Confetti-based analysis, supported by previous studies from other groups, suggest that these differences in plasticity may be driven by spatial cues within the tumour ecosystem [29]. Lineage-related cells belonging to the same clone, depending on their niche, likely experience different environmental signals, such as access to nutrients, interactions with immune and other stromal cells, exposure to growth factors or chemokines, and this probably contributes to their divergent cell fate choices. Interestingly, we identified genes that are associated with clones that contain highly plastic cells, indicating that these cells possess intrinsic properties that guide them through EMT [27, 30, 31]. We report here several markers associated with highly plastic clones, and further studies should molecularly address the relevance of these genes for early stages of tumour development, as well as in tumour invasion.

Furthermore, it is becoming increasingly clear that tumour evolution is not only shaped by competition between clones but also by clonal cooperation. Small clones have been shown to support the growth and expansion of larger clones [32, 33], indicating that cooperation and competition are not mutually exclusive modes of tumour growth [34]. An important question is whether the distinct clones we identified are antagonistic or cooperative. Antagonism may take the form of competition, where clones negatively affect each other, or parasitism, where one clone benefits at the expense of another. Cooperative interactions, in contrast, allow clones to thrive together or benefit mutually. Understanding these dynamics will be crucial to deciphering tumour behaviour and treatment response. A huge gridlock remains in finding a way to isolate individual tumour clones by extracting them from their native tumour environment and probe them in functional assays, without disturbing their physiological behaviour. Interrogating clonal interactions by genetic barcoding should also enable the tracking and retrieval of clones under bottlenecking events such as metastasis and therapy pressure.

In conclusion, by integrating barcode-based lineage tracing with single-cell transcriptomic, we unravelled the transcriptional dynamics underlying both interclonal and intraclonal tumour heterogeneity in breast cancer. This work provides fresh insights into the evolutionary logic of EMT and underscores the importance of integrating lineage information with single cell analyses to disentangle the complexity of tumour heterogeneity. In addition to revealing distinct transcriptional routes encompassing changes in cell state, our approach integrates original techniques, including a probabilistic framework for barcode selection and filtering, a trajectory-informed prognostic analysis of Gene Modules, and a plasticity scoring strategy that links transcriptional heterogeneity to EMT potential and patient prognosis. We believe that these innovative approaches and toolkits will provide a powerful platform for dissecting the interplay between clonal evolution, plasticity, and malignancy in various cancer types.

## Material and Methods

### Mice

ROSACre^*ERT2*^ [35] and N1Cre^*ERT2*^ [36] were crossed with MMTV-PyMT mice [9] and DRAG or Confetti mice [18, 37]. Barcode writing was induced by intraperitoneal injection of tamoxifen dissolved in corn oil (1 mg/10 g of body weight) at 1 month of age. For the analysis of hyperplasia mice were culled at 2 months of age and at 4 months of age for the analysis of late carcinoma.

### Ethics Statement

All studies and procedures involving animals were performed in agreement with the recommendations of the European Community (2010/63/UE) for the Protection of Vertebrate Animals used for Experimental and other Scientific Purposes. Approval was provided by the ethics committee of the French Ministry of Research (reference APAFIS #34364-202112151422480). We comply with internationally established principles of replacement, reduction, and refinement in accordance with the Guide for the Care and Use of Laboratory Animals (NRC 2011). Husbandry, supply of animals, as well as maintenance and care in the Animal Facility of Institut Curie before and during experiments fully satisfied the animals’ needs and welfare. Animal experiments were approved by the ethics committee of the Institut Curie CEEA-IC #118 and by the French Ministry of Research (#55831-2025051216297422 and #34364-202112151422480). All mice were housed and bred in a specific-opportunistic-pathogen-free (SOPF) barrier facility with a 12:12 hours light-dark cycle, with food and water available *ad libitum*. Mice were sacrificed by cervical dislocation.

### Immunofluorescence on Paraffin and Frozen Sections

Tumors were fixed in 4% paraformaldehyde (PFA) at room temperature for 2 hours. For paraffin sections, tumours were embedded in paraffin, sectioned at 7µm, de-paraffinized using a graded alcohol series, and heat-induced antigen retrieval was carried out using antigen retrieval buffer at pH6 or 9.

For cryosections, samples were incubated in 30% sucrose at 4°C for 3 days and then frozen in Tissu-Tek (Sakura). Sections were incubated overnight at 4°C with primary antibodies, followed by incubation with secondary antibodies and DAPI for 2 hours at room temperature.

Primary antibodies used included rabbit anti-Spp1 (22952-1-AP, Proteintech, 1:300), rabbit anti-P-Jun (9164S, Cell Signaling, 1:300), rat anti-K8 (TROMA-1, DSHB, 1:300), rabbit anti-SMA conjugated with AlexaFluor488 (NB600-531, Novus Biologicals, 1:300), chicken anti-K14 (906004, BioLegend, 1:300), rabbit anti-Klf5 (GTX103289, GeneTex, 1:300)and rabbit anti-Hes1 (11988, Cell Signaling, 1:300).

Secondary antibodies used were AlexaFluor488-conjugated anti-chicken IgG (A11039, Invitrogen), Cy3-conjugated anti-rabbit IgG (A10520, Invitrogen), and Cy5-conjugated anti-rat IgG (A10525, Invitrogen).

### Image acquisition

Images were acquired using an LSM780 or LSM880 inverted laser scanning confocal microscope (Carl Zeiss) equipped with 25×/0.8 OIL LD LCI PL APO or 40×/1.3 OIL DICII PL APO objectives. The images were captured with the ZEN Imaging software and processed in Fiji (ImageJ v2.14.0).

### Single-molecule RNA Fluorescence in situ Hybridization (smRNA-FISH)

smRNA-FISH was performed using the RNAscope Multiplex Fluorescent Reagent Kit v2 (Advanced Cell Diagnostics), according to the manufacturer’s instructions. Briefly, paraffin sections were rehydrated with decreasing concentrations of ethanol, pre-treated with the target retrieval reagent (ACD, 322000) for 15 minutes, and then digested with Protease III (ACD, 322381) at 40°C for 15 minutes. Probe hybridization was performed for 2 hours at 40°C. Probes used were mm-Cxcl1-C1 (ACD, 407721) and mm-Lalba-C1 (ACD, 535701). Amplification and binding of dye-labeled probes were then performed. For subsequent immunostaining, sections were incubated in blocking buffer (PBS containing 5% FBS and 2% BSA) for 1 hour. Primary antibodies were incubated overnight at 4°C, followed by 2 hours of incubation with secondary antibodies and DAPI. Slides were mounted in ProLong Diamond Antifade Mountant (Invitrogen-Thermo Fisher Scientific, P36930) for imaging.

### Staining quantification

Nuclei were segmented in Fiji using the DAPI channel. Images were first smoothed using a Gaussian filter to reduce noise, followed by segmentation using Otsu thresholding to generate a binary mask. Segmented nuclei were then reviewed and, if necessary, refined manually or with additional morphological operations to ensure accuracy. Using the same workflow, a mask was generated for the probe channel, with thresholds adjusted according to signal intensity. To quantify probes within each nucleus, regions of interest (ROIs) from the nuclei mask were iteratively selected and analysed. For each ROI, the probe signal was isolated, and particle analysis was performed to summarize probe data per nucleus. For K14- and K17-positive cells, quantification was performed by manual counting of stained cells. Statistical significance for all quantification was evaluated using the Wilcoxon test, with symbols denoting significance levels: * *p* < 0.05, ** *p* < 0.01, *** *p* < 0.001.

### Mammary gland tumour dissociation and single cell sorting

Tumors were harvested and digested with collagenase (Roche, 57981821, 3mg/ml) and hyaluronidase (Sigma, H3884, 200U/ml) for 2 hours at 37°C under agitation. Following washes, cells were incubated with trypsin for 1min, dispase (10888700, Roche, 200U/ml) and DNAseI (D4527, Sigma-Aldrich, 200U/ml) for 5 min and then filtered through a 40 µm cell strainer to obtain a single-cell preparation. Cells were incubated for 45min with the following antibodies (1:100 dilution): APC anti-mouse CD45 (Biolegend), APC anti-mouse Ter119 (Biolegend), APC anti-mouse CD31 (Biolegend), PE anti-mouse Epcam (Biolegend), APC-Cy7 anti-mouse CD49f (Biolegend). Single-cell preparation was resuspended in flow buffer containing PBS, 5 mM EDTA, 1% BSA, 1% FBS, with DAPI for viability staining. Doublets, dead cells (DAPI^*pos*^) and Lin^*pos*^ non-epithelial cells were excluded prior to sorting using FACS ARIA flow cytometer (BD). GFP^*pos*^ cells were collected in BSA 0.04%/PBS. The results were analysed using FlowJo software.

### Barcode amplification on DNA

Sorted cells were lysed in 40µl DirectPCR Lysis Reagent (301-C, Viagen) with 0.4 mg/ml Proteinase K (AM2546,ThermoFisher) and incubated for 1h at 55°C, 30 min at 85°C and 5 min at 94°C. Then, samples were completed up to 130µl with 10mM Tris and shearing was performed for 9 seconds, at a peak power of 70%, with a duty of 20% for 1000 cycles. DNA was split in two replicates before performing PCR. Nested-PCR were finally performed to amplify barcode sequences. 1^*st*^ PCR: 1X of Q5 buffer (B9027S, New England Biolabs), 0.02U/µl of Q5 DNA polymerase, 0.2mM of dNTPs, 0.5µM of each primer for a total of 50µl. PCR program: 98°C for 2min, 98°C for 10sec, 67°C for 10sec, 72°C for 10sec for 30 cycles, 72°C for 5min and hold at 4°C. DNA amplicons were cleaned and size selected using 1.8X SPRI bead-based reagent (B23317, Beckman Coulter) using left side size selection and were eluted in 30µl of RNase/DNase grade H_2_0. 2^*nd*^ PCR: 1X of Q5 buffer, 0.02U/µl of Q5 DNA polymerase, 0.2mM of dNTPs, 0.5µM of each primer for a total of 50µl. Second PCR is designed to add partial Read2 sequence. 2^*nd*^ PCR program: 98°C for 2min, 98°C for 10sec, 67°C for 30sec, 72°C for 30sec for 5 cycles, 72°C for 5min and hold at 4°C. DNA were cleaned and size selected using 1.8X SPRI bead-based reagent using left side size selection and were eluted in 30µl of RNase/DNase grade H20. 3^*rd*^ PCR: 1X of Q5 buffer, 0.02U/µl of Q5 DNA polymerase, 0.2mM of dNTPs, 0.5µM of each primer for a total of 50µl. 3^*rd*^ PCR program: 98°C for 2min, 98°C for 10sec, 67°C for 30sec, 72°C for 30sec for 5cycles, 72°C for 5min and hold at 4°C. Quality and sizes of the DNA products were evaluated using a 4200 TapeStation System (Agilent) with D1000 ScreenTape (Agilent, 5067-5582). Libraries were pooled and cleaned using 1.0X SPRI beads left side selection and eluted in 30µl of RNase/DNase grade H_2_0. Primers used are described in Table 1.

**Table 1.**
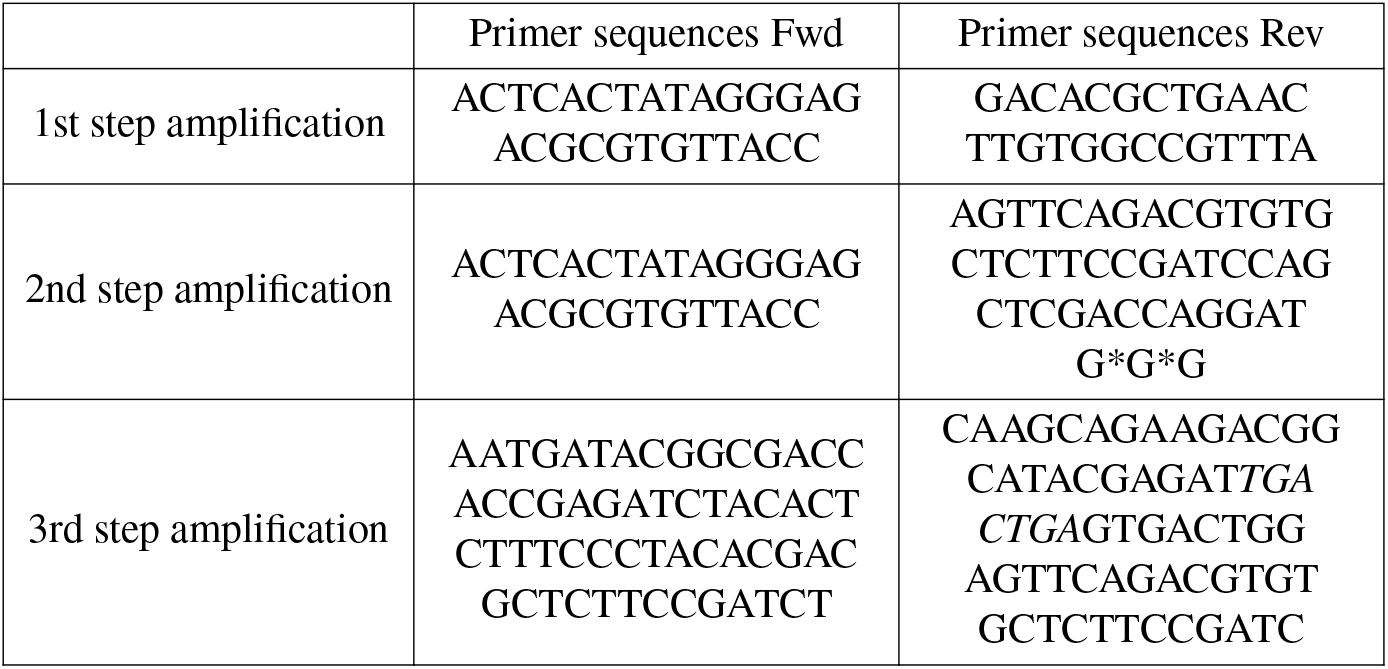
Primer sequences for amplification of barcode sequences on DNA samples.

### Preparation of library for scRNAseq

Single-cell capture and library preparation were performed using the 10x Genomics Chromium Single Cell 3’ v3.1 kit, following the manufacturer’s instructions. The libraries were sequenced on an Illumina NovaSeq 6000 sequencer by the Next Generation Sequencing platform at Institut Curie. Between 6,600 and 16,500 cells were loaded, with a target recovery of 4,000 to 10,000 cells.

### Barcode amplification on amplified cDNA

For the amplification of the barcode sequences, PCR was performed as already described [38], on 10ng of cDNA amplified based on 10x Genomics Chromium Single Cell 3’ v3.1 kit, with 1X of Q5 buffer, 0.02U/µl of Q5 DNA polymerase, 0.2mM of dNTPs, 0.5µM of each primer for a total of 50µl. 1^*st*^ PCR program: 98°C for 3min, 98°C for 15sec, 69°C for 20sec, 72°C for 1min for 15 cycles, 72°C for 1min and hold at 4°C. DNA were cleaned and size selected using 0.6X SPRI bead-based reagent (Beckman Coulter) using left side size selection and were eluted in 40µl of RNase/DNase grade H20. 2^*nd*^ PCR: 1X of Q5 buffer (New England Biolabs), 0.02U/µl of Q5 DNA polymerase, 0.2mM of dNTPs, 0.5µM of each primer, 10µl of DNA for a total of 50µl. 2^*nd*^ PCR program: 98°C for 45sec, 98°C for 20sec, 67°C for 30sec, 72°C for 20sec for 10 cycles, 72°C for 1min and hold at 4°C. DNA were cleaned and size selected using 0.8X SPRI bead-based reagent (Beckman Coulter) using left side size selection and were eluted in 30µl of RNase/DNase grade H20. Quality and sizes of the cDNA products were evaluated using a 4200 TapeStation System (Agilent) with High Sensitivity D5000 ScreenTape (5067-5592, Agilent). Libraries were pooled and cleaned using 1.0X SPRI beads left side selection and eluted in 30µl of RNase/DNase grade H_2_0. Primers used for these PCRs are described in Table 2.

**Table 2.**
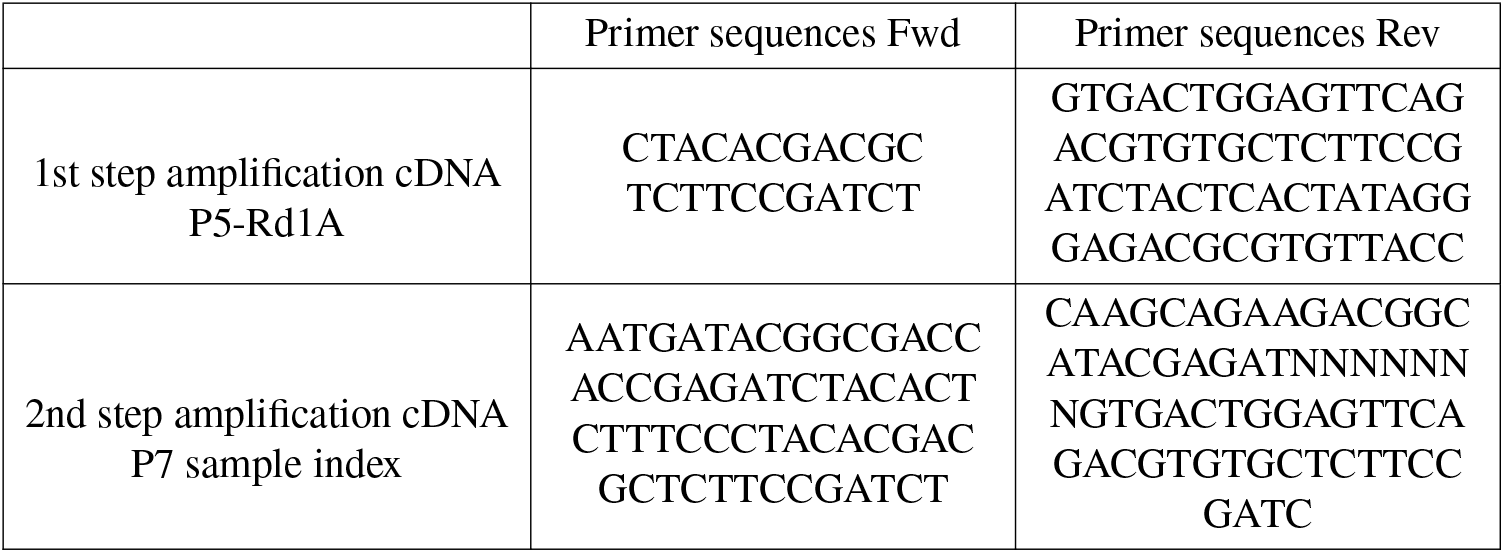
Primer sequences for amplification of barcode sequences on cDNA.

### Barcode sequences extraction

Read1 and Read2 for each sample were concatenated. We used the R package CellBarcode [39], to extract the barcode information from the FASTQ files. The lineage barcode sequences were extracted using the 3^′^ and 5^′^ constant sequences: CGAAGTATCAAG and CCGTAGCAAG corresponding to the V region and J region respectively. For the extraction of barcode sequences from DNA samples, we used the constant sequences CCTCGAGGTCATCGAAGTATCAAG and CCGTAGCAAGCTCGAGAGTAGACCTACT. Sequences covered by less than 2 reads were removed. The dominant barcode of each UMI was extracted only if the number of reads covering this sequence was >75% of total reads for the same UMI. Cells covered by less than 2 UMIs were removed. To be considered valid, the final barcode attributed to a cell must account for at least 55% of the remaining UMIs.

### scRNA sequencing analysis

Raw sequencing reads were processed using Cellranger. The analysis of scRNAseq was performed in R using the Seurat package (v5.2.1) [40, 41, 42]. The following filters were applied to remove doublets and damaged cells: features present in less than 2 cells, cells with less than 500 genes, nCount RNA < 3000 or > 60000, nFeature RNA < 1000 or with a percentage of mitochondrial reads > 20. The data were normalized using the SCTransform function. Principal Component Analysis (PCA) was then performed using the first 15 principal components. Clustering was conducted based on these components using a resolution of 0.4, as determined by the Clustree R package [43], which was used to define the stability of clustering and the appropriate resolution. Uniform Manifold Approximation and Projection (UMAP) was generated using the first 15 PCs, with default parameters. Afterward, integration of all 6 batches was performed using Seurat’s anchor-based workflow (FindIntegrationAnchors and IntegrateData functions). After integration, dataset were scaled and PCA, clustering and UMAP steps were carried out once again on the integrated dataset, using a clustering resolution of 0.4. Markers of the cluster were determined using Seurat’s FindAllMarkers tool. The cell cycle Module Score was calculated using genes from the Gene Ontology term GO0045787 (“positive regulation of cell cycle”) and the EMT score was calculated using the genes associated with GSEA “HALLMARK EPITHELIAL MESENCHYMAL TRANSITION”.

### Barcode filtering

To rigorously filter barcodes and ensure the reliability of clonal assignments, we developed a probabilistic framework that accounts for experimental constraints and is compatible with barcode systems characterized by arbitrary generation probabilities {*q*^*i*^ }_*i*=0,…,*N*_. Each *q*^*i*^ denotes the intrinsic probability of generating barcode *B*^*i*^, as determined by the underlying biological mechanism, in our case, V(D)J recombination. We model barcode generation as a multinomial process in which each of the *n* independent generation events yields one of *N* possible barcodes. The resulting distribution of barcode counts {*g*^*i*^ }_*i*=0,…,*N*_ within a single sample (a single tumour in our case) thus follows a multinomial distribution, and the marginal distribution for a single barcode *B*^*i*^ is binomial: the probability of observing *g*^*i*^ copies of barcode *B*^*i*^ after *n* events is given by *p*^*i*^ (*n, g*^*i*^) = Bin(*n, g*^*i*^).

Barcode generation occurs during the early stages of cancer development, and the total number of generation events *n* is not experimentally accessible. Instead, what can be observed is whether a specific barcode is present in a sample or not. This can be summarised by a new variable *ρ*^*i*^ = min(1, *g*^*i*^). This observation allows us to reframe the problem as estimating the probability that the observed presence of a barcode, *ρ*^*i*^, corresponds to its true generation count, i.e. *g*^*i*^ = *ρ*^*i*^. This essentially means determining whether all cells labelled with *B*^*i*^ originate from a single, unique clone.

To enhance statistical power and robustness, we analysed a large number of cancer samples, each with its own pool of barcodes. The system can be characterized by the generation count matrix 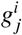, which encodes the number of times barcode *B*^*i*^ was generated in sample *S*_*j*_, and the observation matrix 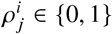, indicating whether *B*^*i*^ was observed in *S*_*j*_. Using this framework, we define the “reliability” of a barcode in a statistical sense, quantified by the probability 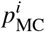 that barcode *B*^*i*^ is monoclonal across all samples. In our analysis, we retained barcodes with 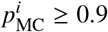, ensuring a high level of reliability.

This approach leverages the fact that all samples are independent realizations of the same multinomial process, sharing the same set of generation probabilities {*q*^*i*^ }_*i*=0,…,*N*_ but differing in the number of generation events *n*_*j*_. This allows us to express the probability of monoclonality as:

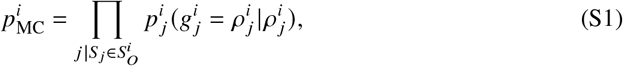

where 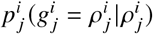 represents the probability that barcode *B*^*i*^ is monoclonal in sample *S*_*j*_ and 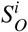 is the set of samples where *B*^*i*^ was observed. Indeed, for barcodes not observed in a certain sample *S*_*j*_, corresponding to 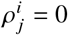, it holds that 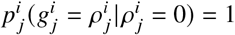. Using Bayes’ theorem and the relationships 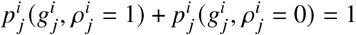 and 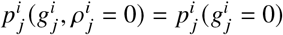, the posterior probability 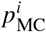 can be expressed as:

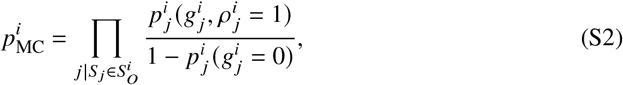

where 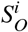 is the set of samples in which *B*^*i*^ was observed.

Since neither the total number of generation events 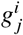 nor the generation probabilities *q*^*i*^ can be directly measured, we approximate these quantities using observable data. The total number of generation events in a sample is approximated as 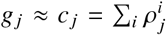, where *c*_*j*_ represents the total number of observed barcodes in sample *S*_*j*_. Similarly, the generation probability *q*^*i*^ is approximated as 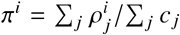, which represents the normalized frequency of barcode *B*^*i*^ across all samples. Substituting these approximations, the posterior probability of monoclonality simplifies to:

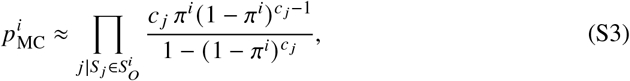

which provides robust estimates for 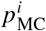.

**Figure M1:**
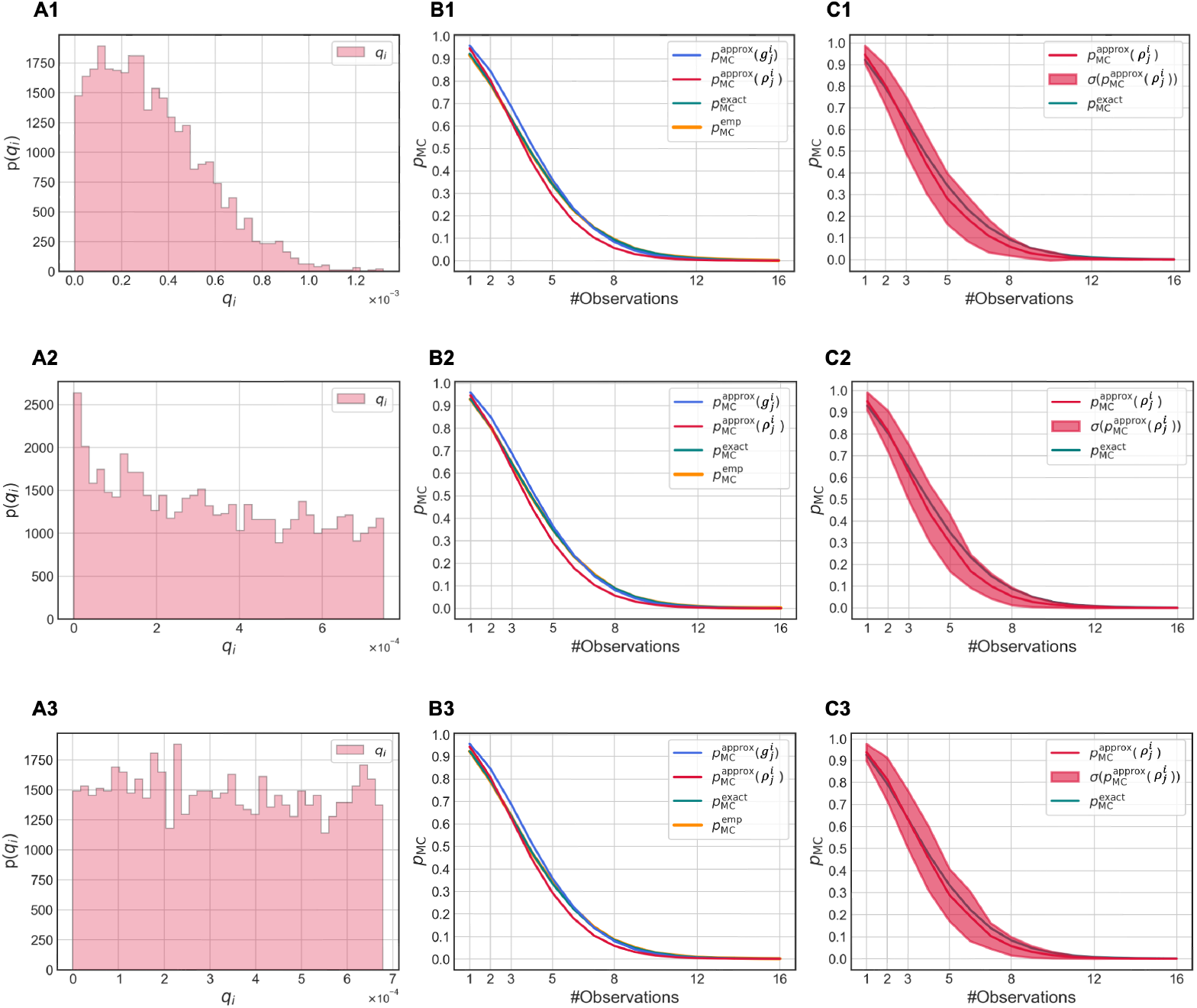
Validation of the barcode monoclonality inference method using numerical simulations. Each row corresponds to a different distribution of barcode generation probabilities *p* (*q*_*i*_): uniform (A), Gaussian (B), and power law (C). **A1, B1** and **C1**. Input distributions used to simulate barcode generation. **A2, B2** and **C2**. Average monoclonality probabilities for barcodes grouped by their total number of observations across samples. The x-axis represents the number of samples in which each barcode was observed. We compared the exact posterior probability 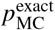 (S2) (green), the approximation (S3) using full generation counts 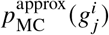 (blue), the approximation (S3) using only presence/absence data 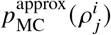 (red), and the empirical monoclonality rate 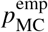 (gold), calculated as the fraction of monoclonal barcodes in each group. **A3, B3** and **C3**. Standard deviation of 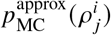 for each observation group, computed within a single simulation run. This variability reflects the noise observed in experimental data (see **Figure M2B**), supporting the realism of the simulations.

Extensive numerical simulations confirmed the robustness of our inference procedure. We tested three different scenarios for the distribution of barcode generation probabilities *p* (*q*_*i*_): a uniform distribution (row A), a Gaussian distribution (row B), and a power-law distribution (row C), as shown in **Figure M1**. For each case, results were averaged over 500 independent simulations. In each row, the first panel shows the predicted distribution of barcode generation probabilities. The second panel displays the average monoclonality probability for barcodes, grouped by how many times they were observed across all samples. Specifically, the x-axis indicates the total number of observations for each barcode (i.e.,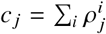), and all barcodes with the same count are grouped together. For each group, we plot the average of three quantities: the approximate posterior probability of monoclonality computed from Eq. (S3) using true generation counts 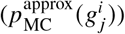, the same approximation using only presence/absence information 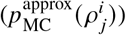, and the exact posterior using Eq. (S2), which relies on full simulation knowledge. We also report the empirical monoclonality probability 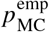, defined as the fraction of barcodes in each group that are truly monoclonal. This curve, shown in gold, perfectly overlaps with the exact analytical prediction. The third panel in each row shows the standard deviation of 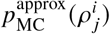 within each group. These fluctuations, calculated from a single simulation (not an ensemble average), closely resemble the variability observed in experimental data (**Figure M2B**). Together, these results confirm that our method reliably estimates barcode monoclonality across a broad range of barcode generation scenarios.

We then applied our filtering procedure to a total of 44 tumour samples: 36 samples for which only barcode-targeted DNA sequencing was performed, and 8 samples with full single-cell transcriptomic data including barcode information. While the 36 bulk-sequenced tumours were not used in downstream analyses, they played a critical role in improving the robustness of the barcode reliability estimates by increasing the total number of barcode observations across samples. For each barcode, we plotted the apparent clonal size, defined as the number of cells sharing a barcode within a single sample, against the number of samples in which that barcode was observed. This number of observations serves as a proxy for the overall prevalence, or “popularity,” of a barcode across the cohort. As shown in **Figure M2A**, the apparent clone size increases exponentially with barcode popularity. Since true clonal size should be independent of barcode frequency, this trend indicates a clear bias that needs to be corrected. We therefore retained only those barcodes with high monoclonality probability, selecting barcodes for which 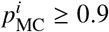. As illustrated in **Figure M2B**, the probability of monoclonality decreases with barcode observation count, and our filtering removes barcodes appearing in more than 5 samples. This yields a set of ~2200 high-confidence barcodes, all over 70% of the original ~3200 barcodes observed across all 44 samples. Among these, about 400 barcodes were detected in the 6 carcinoma samples subjected to scRNA-seq. From this subset, we further selected 43 clones with at least 5 cells each, yielding a total of 482 cells for detailed downstream analysis (**Figure M2C**). This additional threshold on clone size ensures sufficient statistical power for the clonal-level investigations presented in the following sections. Note that, the major reduction in clone number arises not from our monoclonality filtering procedure, but from the requirement on minimum clonal size, which is a downstream analysis constraint. Finally, for the 2 neoplasia samples, we find 254 reliable clones.

**Figure M2:**
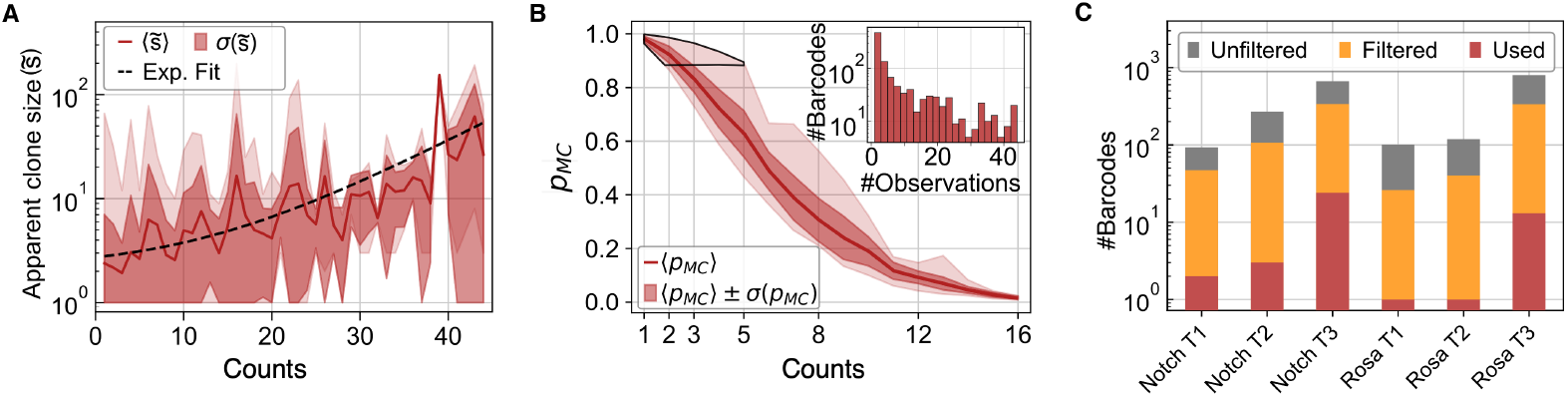
DRAG mouse allows to track hundreds of unique clones in late-stage carcinomas. **A**. Apparent clone size, namely the number of cells with the same barcode in the tumours with single cell transcriptomics, as a function of the number of times that the barcodes appear throughout the whole cohort of tumours. **B**. Probability of barcodes to be monoclonal, i.e. for the apparent clone size to be equal to the real one. As expected, this probability decays monotonically as a function of the number of observations. The inset displays the distribution of the observation number of all barcodes throughout all tumours. **C**. Bar-plot displaying the fraction of filtered barcodes for the 6 tumours with single cell transcriptomics. Orange bars are barcodes for which the probability of being monoclonal is larger than 0.9, whereas the red ones are those for monoclonal barcodes with cell count ≥ 5.

### Fate cluster analysis

To identify groups of clones with similar distributions across transcriptomic space, we represent each clone by a vector *s*, where each component *s*^*i*^ corresponds to the fraction of cells occupying the *i*^th^ cellular state defined in the UMAP plot (**Figure 1A**). This vector captures how each clone is distributed across transcriptional clusters.

To quantify the difference between two clones *a* and *b*, we use the cosine distance:

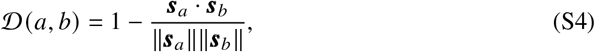

where 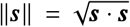 is the Euclidean norm of the vector. This metric captures the angular difference between clones in transcriptomic space and is invariant to their total size. To ensure meaningful comparisons, we applied a size threshold to retain only clones with sufficient numbers of cells, as described in the previous Section. This minimizes noise and improves the robustness of downstream clustering.

We constructed a symmetric distance matrix using the pairwise cosine distances and performed hierarchical clustering using the average linkage method, which iteratively merges clone pairs or groups based on their mean intergroup distance. This analysis revealed seven major clusters of transcriptionally similar clones, which we refer to as *Fate Clusters* (FC0) (**Figure 3A, Figure S4A**). Each Fate Cluster consists of clones that follow comparable transcriptomic trajectories and culminate in similar endpoint cell states. Furthermore, FC0 contains clones that where not assigned to any other Fate Cluster.

### Pseudotime analysis

We next performed pseudotime analysis [20] on Fate Cluster 3 and Fate Cluster 7, which contain the majority of EMT cells. For this analysis, gene expression values were first normalized and corrected for batch effects across samples, and then scaled so that each gene had a mean of zero and a variance of one. Pseudotime analysis provides a way to reconstruct cellular trajectories by arranging cells along a continuous axis of transcriptional progression, thus transforming static single-cell data into a dynamic representation of state transitions. This approach enabled us to infer how cells gradually change their transcriptional programs, culminating in the EMT phenotype. Importantly, the endpoint of the trajectory was not manually imposed but consistently emerged at the EMT-enriched region of transcriptomic space (**Figure 3C-D**).

For Fate Cluster 3, the pseudotime origin was initialized using the cell in the alveolar-enriched cluster with the highest expression of *Lalba*, a hallmark gene of alveolar differentiation. For Fate Cluster 7, the starting point was set as the cells in the AP-1 activated cluster with the highest expression of *Krt8*, a canonical luminal marker. To compute pseudotime, we constructed a diffusion map representation of the transcriptomes and derived diffusion pseudotime (DPT) values by initializing trajectories from the selected starting cells. For Fate Cluster 3, we chose 10 diffusion components and 15 neighbors, while for Fate Cluster 7, we chose 5 diffusion components and 5 neighbors. To display gene expression trends along pseudotime, we further applied smoothing using generalized additive models (GAMs), which provided a clear visualization of gradual transcriptional changes across each trajectory.

**Figure 3E-G**, shows gene expression in relation to pseudotime values, also presents batch-corrected and scaled data.

### Gene modules using Hotspot

Following the identification of transcriptional trajectories using pseudotime analysis, we sought to uncover gene programs driving these transitions. To do so, we applied the *Hotspot* algorithm to identify gene modules with coherent co-expression patterns across the cell state space. As required by Hotspot, we provided a tree structure capturing relationships between similar clones, derived from the hierarchical clustering shown in **Figure 3A**. We used raw count data and fitted gene–cell relationships with a depth-adjusted negative binomial (DANB) model, which provides a baseline for testing local co-expression. To account for stochastic variability, we collected many independent realizations and aggregated them through consensus clustering. The consensus matrix was then evaluated using silhouette metrics together with a z-score filter (*p* < 0.01) to identify and reassign genes that were inconsistently grouped. We further explored different neighborhood sizes and gene thresholds, comparing results across parameter choices, and found that a neighborhood size of 30 and a minimum gene threshold of 25 provided a reasonable balance. Under these settings, we obtained 14 gene modules, comprising 2422 genes, that reflect functionally relevant programs involved in cancer evolution (**Figure 4A**).

To quantify the activity of each gene module across Fate Clusters, we calculated a score for every gene module in each cell. This was done by computing the average expression of the genes belonging to a given module and comparing it to the average expression of a control set of genes of the same size, randomly selected from the transcriptome. The resulting score reflects the relative activation of the module, standardized with respect to a reference distribution. For this analysis we used batch-corrected and scaled data, so that differences in module activity would primarily reflect biological variability rather than technical effects such as sequencing depth or inter-sample batch differences. Higher scores indicate stronger relative activity of the module in that cell, enabling association of specific gene programs with distinct cellular states or evolutionary trajectories. To summarize and compare module activation across Fate Clusters, we computed the average Gene Module Score within each Fate Cluster. These average values were then visualized in a heatmap (**Figure 4C**), revealing which gene modules are selectively enriched along each Fate Cluster.

### Prognostic Analysis Using Human Cancer Datasets

To link our findings from the mouse model to clinical data and assess their potential translational relevance, we evaluated whether the gene modules identified in clonal trajectories are associated with patient prognosis. This analysis aimed to determine whether transcriptional programs active in specific evolutionary routes correlate with survival outcomes in human cancers.

We first mapped the mouse genes in each module to their human orthologs. Using these mapped gene sets, we computed Module Scores for a cohort of 1,094 breast cancer (BRCA) patients based on bulk RNA-sequencing data from TCGA-BRCA [22]. The data were treated analogously to single-cell observations, with each patient considered as one unit. Expression values were normalized and subsequently log-transformed log(1 + *x*) before scoring. For each module, patients were stratified into two groups depending on whether their score was above or below the cohort mean. The mean threshold was chosen to highlight patients with relatively high module activity, as these cases were of particular interest for prognostic evaluation.

To assess the relationship between module activity and clinical outcomes, we compared survival distributions between the two groups using the log-rank test. This test measures whether differences in survival times are statistically significant based on observed versus expected events. Multiple hypothesis testing was addressed by applying the Benjamini-Hochberg criterion, which controls the false discovery rate. Results for the gene modules that passed this significance threshold are shown in **Figure 4**.

This approach enabled us to associate gene modules, originally identified in mouse clonal trajectories, with patient prognosis, offering a bridge between tumour evolution in experimental models and clinical outcomes.

### Plasticity score

To quantify plasticity within each barcoded clone, we developed a score that combines both the number of transcriptional states represented within a clone and their progression along pseudotime. First, cells were ordered by pseudotime within their respective clone, starting from the earliest cell, which was assigned a plasticity score of zero. As we progressed along pseudotime, the score was incremented whenever a cell occupied a transcriptional state (as defined by the UMAP clusters in **Figure 1A**) not previously encountered within that clone. This procedure ensures that diversity in cell states contributes to the score only once, when first observed. For each of the 282 analysed cells, the final plasticity score was then computed as the product of this cumulative state count and the cell’s pseudotime value. In this formulation, pseudotime acts as a proxy for differentiation potential, so cells that appear later and diversify into novel states receive proportionally higher scores. Consequently, clones that spread across multiple transcriptional states and extend further along pseudotime attain higher scores, reflecting broader transcriptional repertoires and evolutionary progression. Conversely, clones confined to a single state accumulate low scores despite their size. This design has two advantages: it highlights cells within large, diverse clones that are more likely to adapt to changing conditions, while also penalizing cells that remain in limited transcriptional niches. High-plasticity cells were defined as the top 10% of the distribution of scores across all cells, and low-plasticity cells as the remainder.

To probe the molecular basis of plasticity, we compared gene expression profiles between high- and low-plasticity cells within each Fate Cluster. Differentially expressed genes were identified by retaining only those with an adjusted p-value below 0.01 and an absolute log_2_ fold change greater than 2, ensuring both statistical significance and strong effect size. This analysis revealed transcriptional programs specifically enriched in highly plastic cells. We also contrasted low-plasticity cells originating from clones that contained high-plasticity cells with those from clones entirely composed of low-plasticity cells. Interestingly, even among low-plasticity cells, those belonging to heterogeneous clones exhibited gene expression patterns indicative of a primed state, suggesting that plasticity can be encoded as a broader clonal property rather than emerging only at the level of individual cells.

### Gene Ontology Enrichment Analysis

To functionally annotate gene modules, Gene Ontology (GO) enrichment analysis was performed using the R package clusterProfiler (v4.10.1) with org.Mm.eg.db (Mus musculus annotation database, v3.18.0). Gene symbols associated with each module were extracted and analysed separately. For each module, GO terms were filtered to retain only those with an adjusted p-value < 0.05 and a minimum gene count of three. The top five enriched terms per module, ranked by adjusted p-value, were selected for visualization. Bar plots were generated using ggplot2 (v3.5.2).

## Data availability

10X scRNA-sequencing data generated in this study is accessible on the Gene Expression Omnibus GEO repository (GSE292488).

## Acknowledgments

We wish to warmly acknowledge the flow cytometry and cell sorting platform at Institut Curie for their technical support, the in vivo experimental facility for help in the maintenance and care of our mouse colony, and the Cell and Tissue Imaging Platform PICT-IBiSA at Institut Curie (member of the French National Research Infrastructure France-Bioimaging, ANR-10-INBS-04) for their expertise and assistance. High-throughput sequencing was performed by the ICGex NGS platform of the Institut Curie supported by the grants ANR-10-EQPX-03 (Equipex) and ANR-10-INBS-09-08 (France Génomique Consortium) from the Agence Nationale de la Recherche (“Investissements d’Avenir” program), by the ITMO-Cancer Aviesan (Plan Cancer III) and by the SiRIC-Curie program (SiRIC Grant INCa-DGOS-465andINCa-DGOS-Inserm-12554). Data management, quality control and primary analysis were performed by the Bioinformatics plat-form of the Institut Curie. We are very grateful to all members of the Fre laboratory for support, critical reading of the manuscript and constructive discussions. This work was funded by Paris Sciences et Lettres (PSL* Research University) (grant C19-64-2019-228), the French National Research Agency (ANR) grant numbers ANR-21-CE13-0047 and ANR-22-CE13-0009, the Medical Research Foundation FRM “FRM Equipes” EQU201903007821, the FSER (Fondation Schlumberger pour l’éducation et la recherche) FSER20200211117, the Association for Research against Cancer (ARC) label ARCPGA2021120004232 4874, the Worldwide Cancer Research Foundation 24-0216, GEFLUC “les Entreprises contre le Cancer” and by Labex DEEP ANR-Number 11-LBX-0044 to SF. C.M. was funded by a postdoctoral fellowship from ARC (ARCPDF12021020003033).

The funders had no role in study design, data collection and analysis, decision to publish, or preparation of the manuscript.

## Author contributions

C.M. conceptualized the project, designed experiments and performed experimental work. I.D.T. developed the mathematical models and conducted computational analyses. M.H. and D.W. contributed to experimental work, and W.S. contributed to the initial analyses. C.C. and L.P. provided the primer sequences and protocol for amplifying the transcribed barcodes. S.F. and S.R. supervised and provided funding for the project. C.M, I.T., S.R. and S.F. wrote the manuscript.

