## Supplementary figures for "Genetic barcoding of individual cells links cancer evolutionary trajectories and prognostic outcomes"

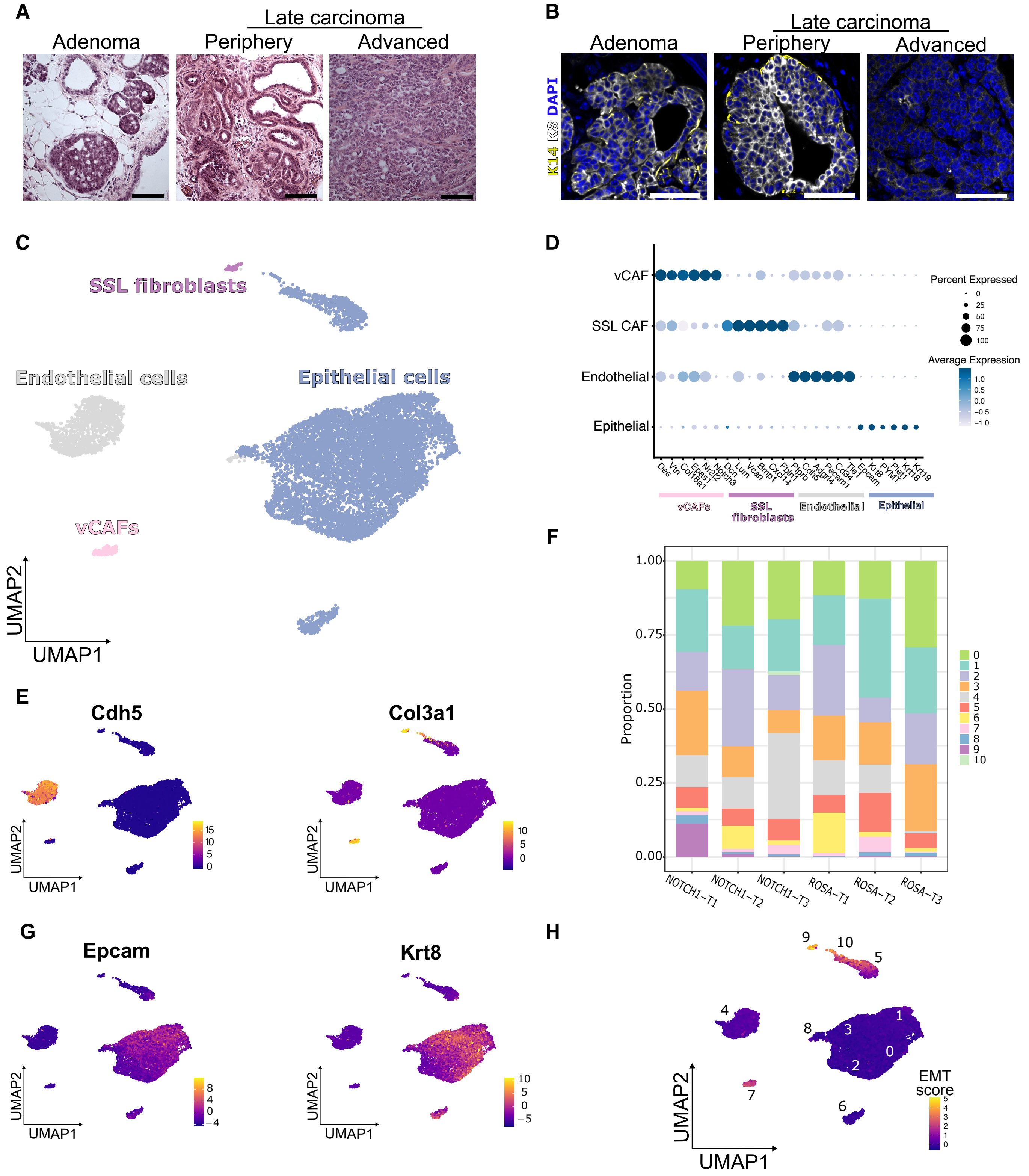


**Figure S1: Identification of the different populations of cells coexisting in MMTV-PyMT late-stage carcinomas.**

**Figure S1:** **A.** Representative Hematoxylin and Eosin staining of adenoma and late-stage carcinoma sections from MMTV-PyMT tumours. Scale bars represent 100 µm. **B.** Immunostaining of tumour sections at different stages using luminal (K8) and basal (K14) markers. Scale bars represent 50 µm. **C.** UMAP indicating the 4 distinct cell types identified in 6 late carcinomas. Each colour represents a cell type. vCAFs denote vascular CAFs and SSLs are associated with Steady State-Like CAFs [44, 12]. **D.** Dot plot displaying the expression of markers associated with the different cell types. Dot size represents the percentage of cells expressing the gene gene, and the colour (white to dark blue) represents average expression. **E.** UMAP showing the expression of *Cdh5*, specific of endothelial cells, and *Col3a1* specific to fibroblasts. **F.** Barplot indicating the proportion of each cluster (different colours) within one late carcinoma. **G.** UMAP showing expression of markers specific to luminal cells. **H.** UMAP showing the EMT score based on genes belonging to the hallmark "epithelial to mesenchymal transition".

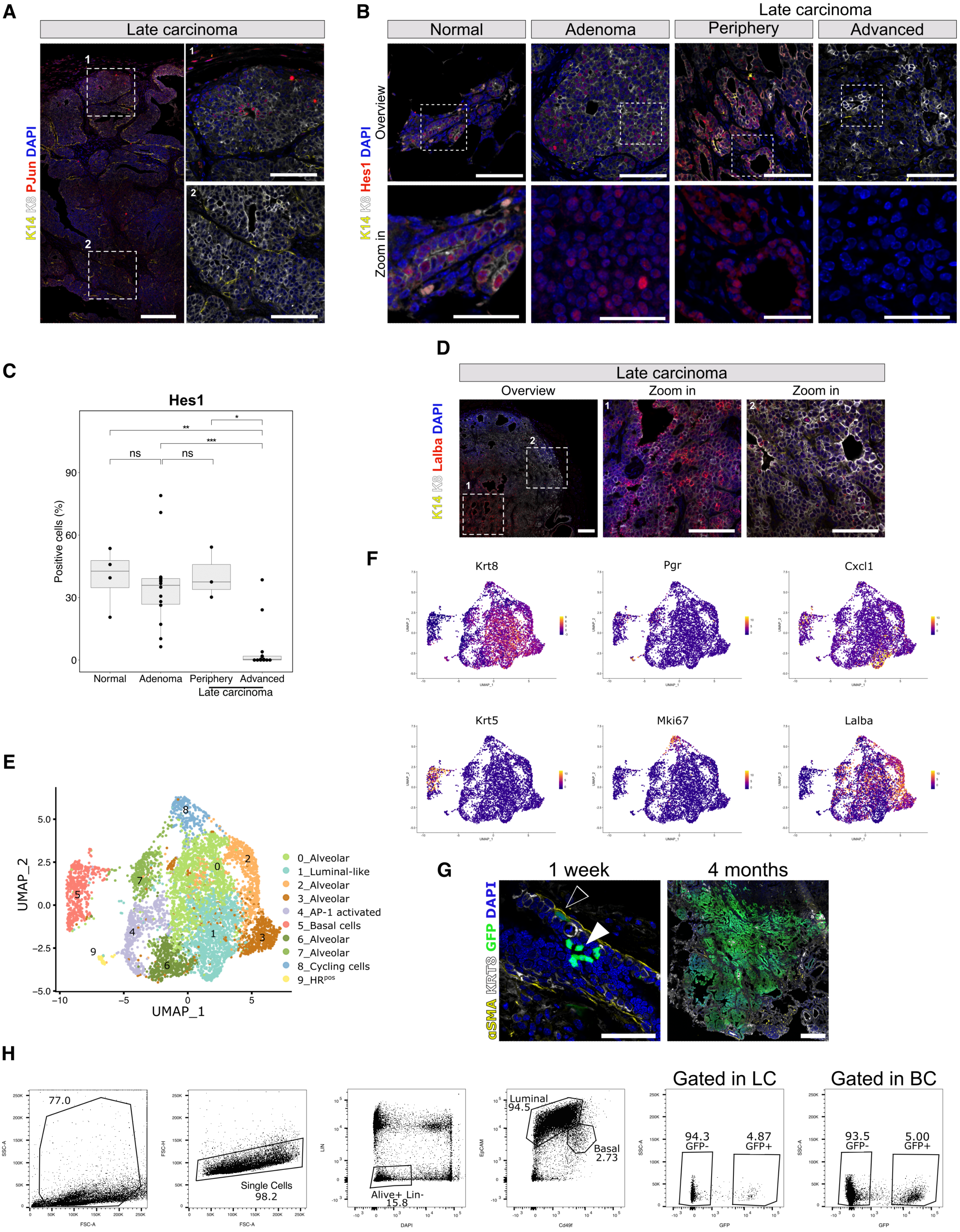


**Figure S2: Identification of the different populations of cells coexisting in MMTV-PyMT late-stage carcinomas.**

**Figure S2:** **A.** Immunofluorescence of representative late carcinoma sections for PJun illustrating the heterogeneity of the expression inside a tumour. 1 indicates the tumour periphery and 2 the tumour centre. **B.** Immunofluorescence for Hes1 at different stages of tumour progression. Scale bars represent 50 µm for the overview and 25 µm for the zoom-in. **C.** Quantification of Hes1-positive cells across different stage of tumour progression. **D.** smRNA FISH of representative late carcinoma tumours for the marker *Lalba* showing heterogeneous expression at the late carcinoma stage. **E.** UMAP plot showing the different clusters identified after integration of 2 individually sequenced adenomas. **F.** UMAP plots showing expression of markers used to identified clusters defining late carcinomas tumours. **G.** Immunostaining of the mammary gland showing GFP$^{pos}$ cells in both luminal and basal compartments after a 1-week chase, and clonal expansion of a GFP$^{pos}$ clone after a 4-month chase. Left panel: scale bar = 50 µm; right panel: scale bar = 200 µm. **H.** Gating strategy for single-cell sorting and selection of barcoded (GFP$^{pos}$) cells.

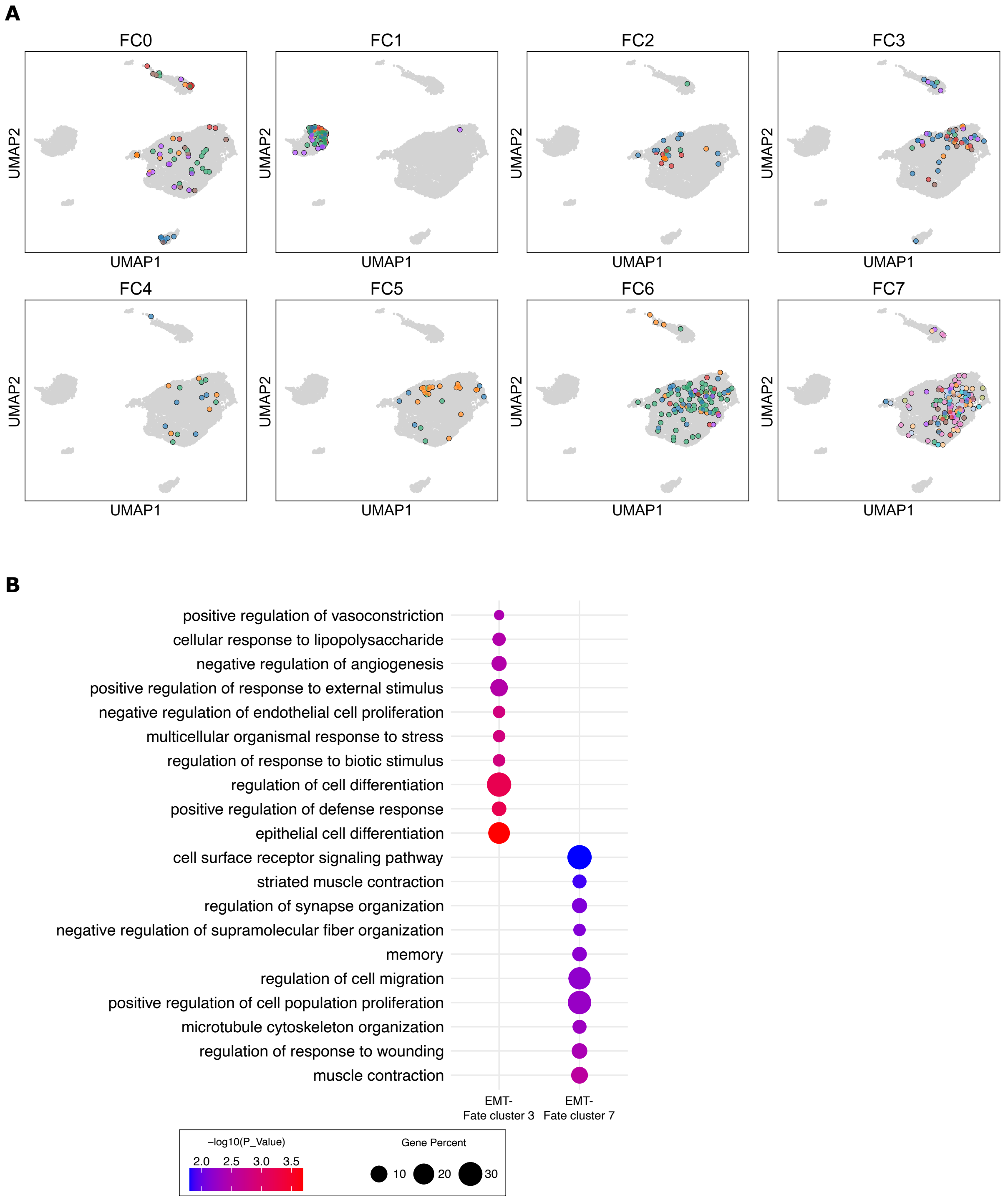


**Figure S3: Clonal composition and characterization of Fate Clusters. A.** Clonal composition of all Fate Clusters. Each colour represents a distinct clone. **B.** Gene Ontology (GO) terms associated with genes from the EMT cluster, derived from Fate Cluster 3 or Fate Cluster 7.

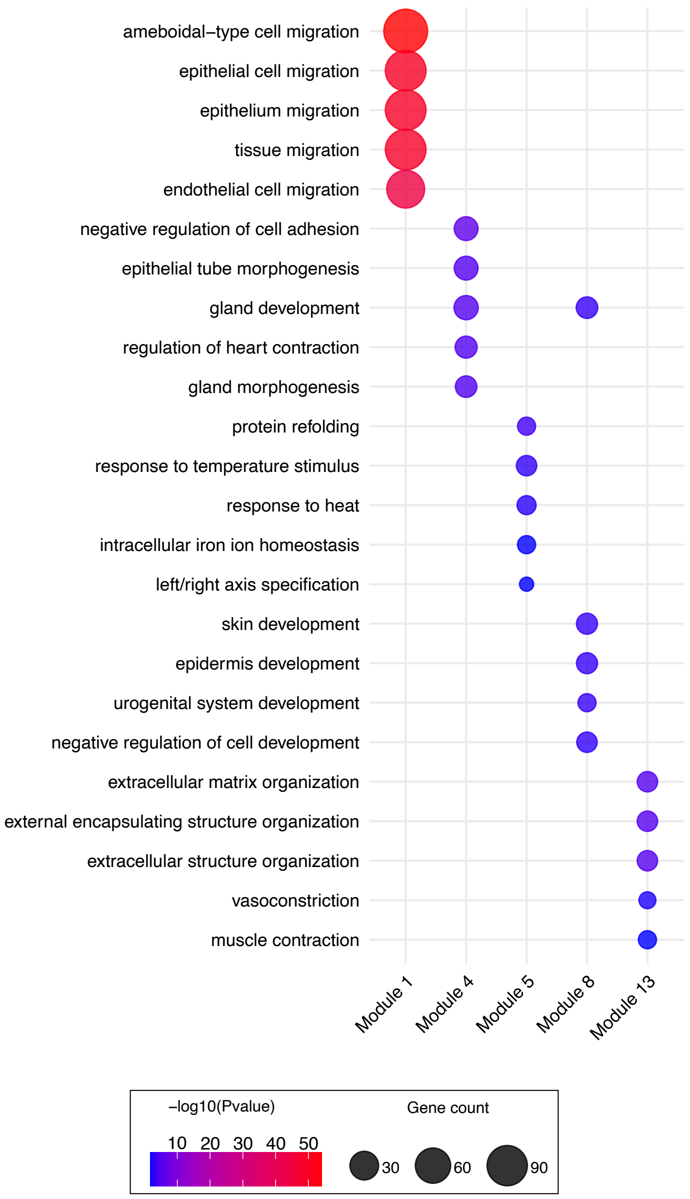


**Figure S4: Biological processes associated with defined gene modules.** GO terms associated with specific modules.

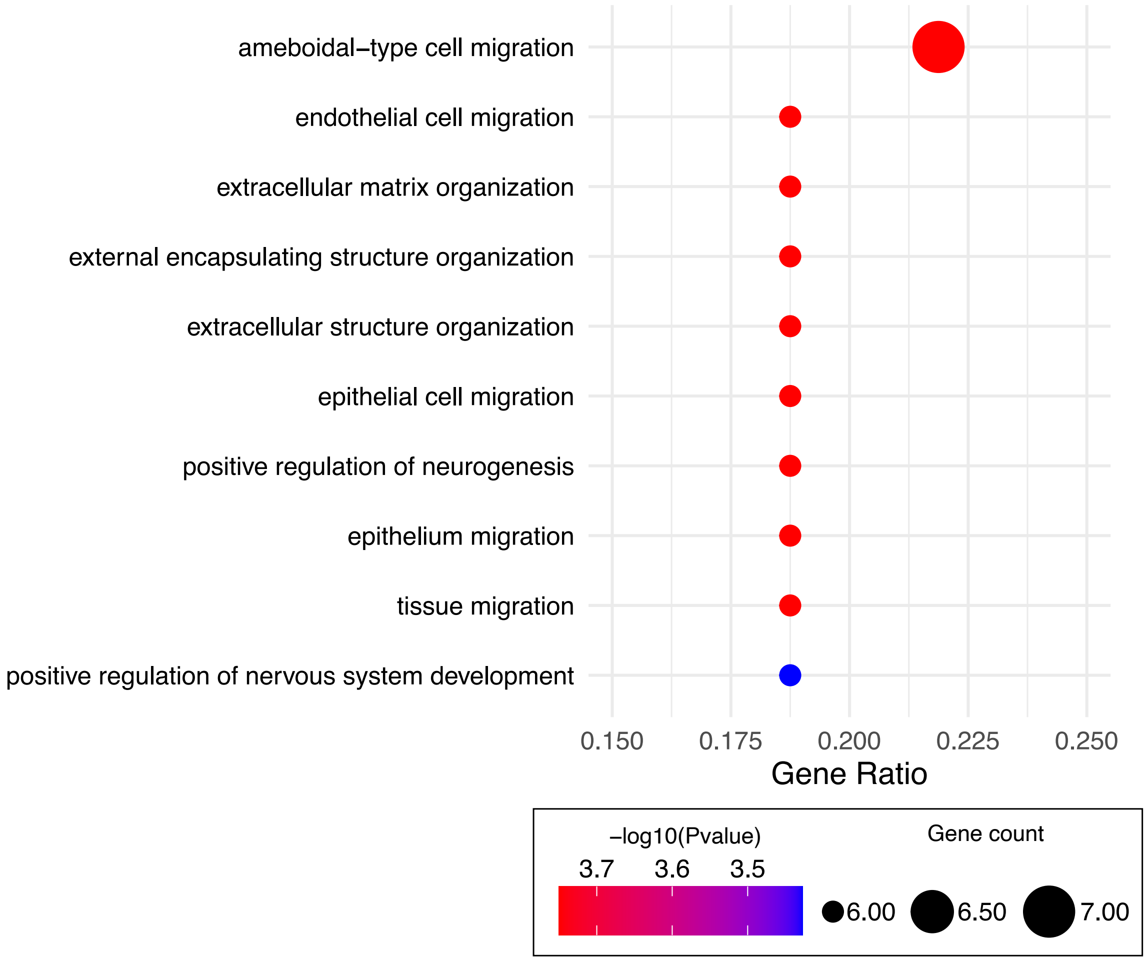


**Figure S5: GO terms associated with the genes expressed by highly-plastic cells**
